# IL-1β/IL-6 signaling circuit in the tumor microenvironment drives prostate cancer development

**DOI:** 10.64898/2026.07.30.741572

**Authors:** Young Sun Lee, Jimmy L. Zhao, Agnieszka Chryplewicz, Max Land, Roshan Sharma, Joseph Chan, Perianne Smith, Sanjay Kottapalli, Linda Fong, Zhenghao Chen, Eric Bent, HuiYong Zhao, Elisa de Stanchina, Wenfei Kang, Shevin Narine, Eric Rosiek, Ning Fan, Kayla Lawrence, Erolcan Sayar, Anuradha Gopalan, Ojasvi Chaudhary, Tianhao Xu, Ignas Masilionis, Ronan Chaligne, Michael Haffner, Rodrigo Romero, Dana Pe’er, Brett S. Carver, Charles L. Sawyers

**Affiliations:** Human Oncology and Pathogenesis Program, Memorial Sloan Kettering Cancer Center, New York, NY 10065, USA; Computational & Systems Biology Program, Sloan Kettering Institute, Memorial Sloan Kettering Cancer Center, New York, NY 10065, USA; Calico Life Sciences LLC, San Francisco, CA 94080, USA; Antitumor Assessment Core Facility, Memorial Sloan Kettering Cancer Center, New York, NY 10065, USA; Molecular Cytology Core Facility, Memorial Sloan Kettering Cancer Center, New York, NY 10065, USA; Department of Genitourinary Oncology, Memorial Sloan Kettering Cancer Center, New York, NY 10065, USA; Division of Human Biology, Fred Hutchinson Cancer Center, Seattle, WA 98109, USA; Department of Pathology, Memorial Sloan Kettering Cancer Center, New York, NY 10065, USA; Department of Laboratory Medicine and Pathology, University of Washington, Seattle, WA 98195, USA; Department of Surgery, Memorial Sloan Kettering Cancer Center, New York, NY 10065, USA; Howard Hughes Medical Institute, Chevy Chase, MD 20815, USA

## Abstract

Despite considerable progress in elucidating mechanisms leading to castration-resistant prostate cancer (CRPC), insight into the early stages of prostate cancer initiation and progression remains limited. Genomic drivers of prostate cancer initiation have been defined through patient tumor sequencing, but the subsequent events responsible for local tissue invasion are poorly understood. Here we leverage a well-studied genetically engineered mouse prostate cancer model (Hi-Myc) that, based on robust and reproducible kinetics for transitioning from pre-invasive prostatic intraepithelial neoplasia (PIN) to invasive prostate adenocarcinoma (PCa), provides an ideal system to systematically address this question using single-cell analysis. Surprisingly, the transcriptomic profiles of early PIN lesions are indistinguishable from those of late-stage, highly invasive tumor cells, suggesting that MYC activation at the PIN stage establishes a transcriptional program that is fully capable of driving invasion but is restrained by the local tumor microenvironment (TME). Indeed, we find that progression to PCa is associated with progressive infiltration of IL-1β^+^ tumor-infiltrating macrophages at the PIN stage that, based on immunodepletion and cytokine neutralization experiments, are required for the PIN-to-PCa transition. Mechanistically, IL-1β from macrophages acts directly on prostate fibroblasts, leading to the release of IL-6, which drives invasion by activating IL-6R in tumor cells. Collectively, these findings identify a pro-tumorigenic IL-1β/IL-6 signaling circuit mediated through local macrophages and fibroblasts that unleashes the full oncogenic potential of a cancer driver (MYC) activated at the PIN stage. We also find evidence of this circuit in other (non-MYC-driven) prostate cancer models as well as human prostate and lung adenocarcinoma, with implications for TME-specific targeted therapeutics in early-stage disease.

## Introduction

Extensive work on castration-resistant prostate cancer (CRPC) progression has revealed both cancer cell-intrinsic and -extrinsic changes that sustain cancer cell survival in advanced disease.^1–12^ Accumulation of second hit genomic alterations, such as *AR* amplification and loss of *Rb1*, promotes continued tumor growth and drives disease progression through mechanisms including lineage plasticity and transformation into neuroendocrine prostate cancer.^1,10,11^ Within the TME, fibroblasts and immunosuppressive neutrophils drive resistance to androgen deprivation therapy through cytokine release.^12,13^ Fibroblasts, as a primary source of fibroblast growth factor (FGF), can also activate JAK/STAT signaling in cancer cells, a step required for the transition to neuroendocrine disease.^11^

In contrast, mechanisms governing the earliest stages of prostate adenocarcinoma (PCa) development remain poorly understood. Genomic sequencing of patient samples has identified common oncogenic drivers^14–18^ of PCa—*ERG, FOXA1*, *SPOP*, and *MYC*—but the stepwise iteration of genetic and microenvironmental changes that enable pre-cancerous lesions to progress to invasive adenocarcinoma has not been fully defined. This gap in knowledge stands in contrast to other types of adenocarcinomas with well-defined progression sequences, such as *Kras* mutation followed by *Tp53* alteration in pancreatic cancer,^19–22^ or *Apc* deletion followed by *Kras, Smad4*, or *Trp53* alterations in colorectal cancer.^23,24^ Consequently, whether progression from PIN to invasive PCa reflects additional changes within tumor cells or depends on remodeling of the TME remains unknown.

The role of inflammation in early tumorigenesis, well-established in several epithelial malignancies,^25–32^ raises the question of whether similar mechanisms drive PCa development. In pancreatic and lung cancer, inflammation induces plasticity or wound-healing programs that promote malignant transformation.^33–36^ In the prostate, single-cell analysis of human tumor samples across disease progression revealed infiltration of distinct immune populations at different stages.^37–40^ In addition, prostatic inflammatory atrophy (PIA), lesions marked by high neutrophil infiltration and atrophic epithelium, is associated with cancer initiation, implicating immune cells in early disease.^41^ However, the mechanistic accounting of how TME changes promote progression from premalignancy to invasive PCa has not been established.

Clinical evidence supporting a functional role for the TME during tumorigenesis emerged unexpectedly from the CANTOS trial of over 10,000 patients, in which treatment with the IL-1β-neutralizing antibody canakinumab reduced overall cancer incidence relative to the control group. IL-1β is produced by myeloid cells, including macrophages, monocytes, and neutrophils, and mediates canonical inflammatory IL-1 signaling.^42,43^ Subsequent CANOPY trials attempted to leverage this finding in patients with established non-small cell lung cancer but failed to demonstrate clinical benefit of canakinumab,^44–47^ suggesting that the TME exerts effects on tumor initiation/progression in a stage-specific manner. The divergent clinical contexts—the CANTOS trial addressing cancer incidence and CANOPY trials addressing progression of established disease—underscore the importance of understanding TME contributions early during the transition from premalignancy to invasive cancer.

Here, we used the Hi-Myc prostate cancer model to investigate the mechanisms by which early changes within tumors drive cancer development. In this widely used transgenic model, tumorigenesis is driven by overexpression of a single oncogene, *MYC*, the most commonly amplified gene in human cancer, with expression restricted to prostatic epithelial cells by the probasin promoter.^48–57^ Without requiring an additional genetic hit, Hi-Myc tumors progress gradually from pre-cancerous PIN to invasive PCa over months, closely recapitulating the histological evolution of human prostate cancer. PIN lesions arise as early as 2 weeks of age, with invasive features emerging around 6 months and fully penetrant PCa by 8 months. This defined, reproducible disease kinetics provides a powerful platform for dissecting cellular and molecular events that occur during the transition from premalignancy to invasive cancer.

Using single-cell RNA-sequencing (scRNA-seq), we profiled transcriptional changes across all cell types during the PIN-to-PCa transition. Strikingly, *MYC* transgene-expressing cells were already transcriptionally poised for invasion as early as 6 weeks, with little further change in their intrinsic program as disease progressed, suggesting that TME remodeling gates the transition to malignancy. In contrast to the stable transcriptional programs within tumor cells, the TME underwent substantial changes, particularly within macrophage and fibroblast populations. Functional studies using antibody blockade and mouse organoids identified a pro-inflammatory IL-1β/IL-6 signaling circuit, in which macrophage-derived IL-1β activates prostate fibroblasts to upregulate IL-6, thereby promoting tumor progression. Dual blockade of IL-1β and IL-6 signaling was more effective than single-agent IL-1β inhibition at delaying tumor progression. Analysis of patient datasets provided evidence that this inflammatory signaling axis is conserved across prostate and lung adenocarcinomas, suggesting that macrophage-fibroblast crosstalk represents a broadly shared mechanism regulating the transition from premalignancy to invasive cancer.

## Results

### The transcriptional program of preneoplastic PIN lesions is indistinguishable from invasive PCa tumor cells

The Hi-Myc model faithfully recapitulates the morphologic progression observed during human prostate tumorigenesis, characterized by prominent nucleoli in PIN lesions and subsequent loss of the basal cell layer as tumor cells invade the surrounding stroma.^58^ To confirm the kinetics of disease progression previously reported in the Hi-Myc model, we collected whole prostates from mice at ages 6 weeks, 3 months, 6 months, and 8 months, and performed histology and immunohistochemistry (IHC) for transgene expression (TG-MYC), proliferation (Ki67), and basement membrane integrity (SMA). As expected, highly proliferative PIN lesions were detected at 6 weeks and persisted through 6 months, with a doubling of prostate weight due to an increase in PIN volume (**Fig 1A, S1A, S1B**). At 8 months, nearly all mice had evidence of invasive PCa, highlighting the 6 to 8 month interval as the critical period for the PIN-to-PCa transition. Having validated the timing of this transition, we then performed scRNA-seq on age-matched whole prostates from wild-type and Hi-Myc mice across these same timepoints (6 weeks, 3 months, 6 months, 8 months) to examine tumor cell-intrinsic and- extrinsic changes (**Fig 1B-C, S1C-D**).

**Figure 1.**
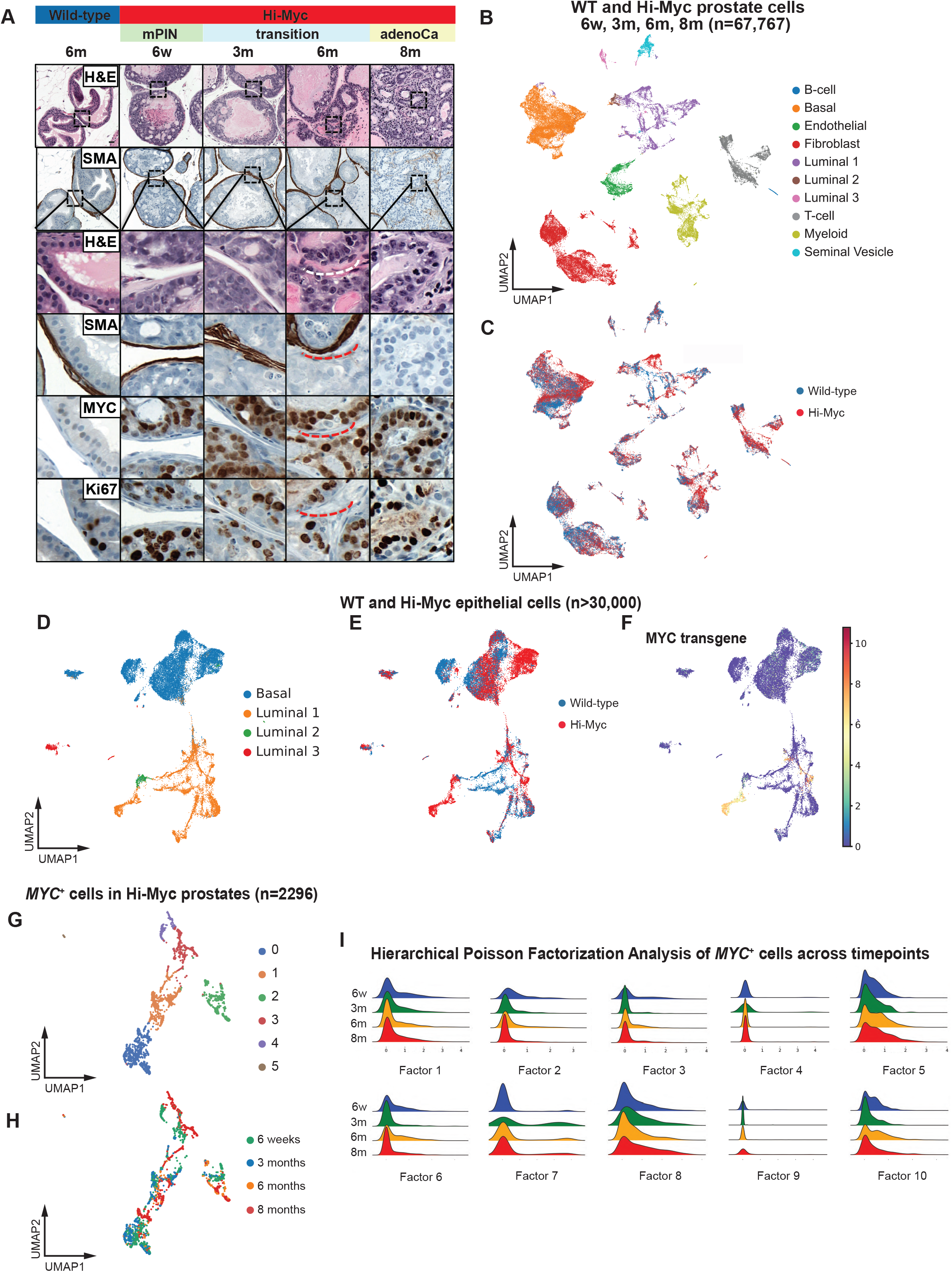
Tumor-intrinsic transcriptional programs remain stable during the PIN-to-PCa transition. (A) Representative H&E and IHC images of wild-type and Hi-Myc prostates at multiple timepoints, shown at low (40X) and high (200X) magnification. Scale bar: black 20 µm, white 5 µm. (B) UMAP of cell types identified by scRNA-seq. Data were pooled from wild-type and Hi-Myc prostates at 6 weeks, 3 months, 6 months, and 8 months of age (n = 67,767). (C) UMAP showing genotypes across cell types. (D) UMAP of epithelial cells in wild-type and Hi-Myc prostate cells (n > 30,000). (E) UMAP displaying genotypes of epithelial cells. (F) UMAP of human *MYC* transgene expression in wild-type and Hi-Myc epithelial cells. (G) UMAP of Phenograph clusters in *MYC*^+^ epithelial cells from Hi-Myc prostates (n = 2,296). (H) UMAP representation of cells from different timepoints during tumor progression. (I) scHPF analysis of *MYC*^+^ cells across timepoints for *de novo* identification of latent gene expression programs.

Focusing first on epithelial cells (*Epcam*^+^), we identified all previously reported major epithelial cell populations (Luminal_1, 2, 3, Basal). As expected, we observed clear separation of *TG-MYC*^+^ cells from wild-type epithelial cells (**Fig 1D-F**).^59^ Gene set enrichment analysis (GSEA) between *TG-MYC*^+^ and wild-type luminal cells demonstrated upregulation of MYC and cell proliferation pathways and downregulation of cell junction and MYC-repressed pathways (**Fig S1E**). *TG-MYC^+^* cells predominantly expressed a Luminal_1 (L1) lineage signature throughout the PIN-to-PCa transition, including markers such as *Prom1, Pbsn, Hoxb13, Nkx3.1* (**Fig S1F-G**). The persistent L1 identity of *TG-MYC*^+^ cells was confirmed by flow cytometry at both PIN (6-month-old) and PCa (12-month-old) stages (**Fig S1H**). Although *TG-MYC^+^* cells are transcriptionally closest to L1 cells, they also expressed stem and progenitor cell markers (*Sca-1, Trop2, CD44*) seen in Luminal_2 (L2) cells in wild-type prostates, a finding confirmed at the protein level by flow cytometry (**Fig S1I-J**). Spatially, we observed clear anatomic separation of TROP2^+^ (L2 marker) and PROM1^+^ (L1 marker) in the proximal and distal regions of prostate lobes from wild-type mice, as reported previously.^59,60^ In contrast, analogous sections from Hi-Myc mice showed expansion of double positive (PROM1^+^/TROP2^+^) cells both proximally and distally (**Fig S1K**). Thus, *TG-MYC^+^* expression drives expansion of a predominantly L1-like epithelial population, with co-expression of L2 markers associated with stemness. Of note, this transcriptional phenotype differs from the predominantly L2-like signature seen in PCa models initiated by other common driver alterations such as *ERG* and *PTEN*.^61,62^

Given the histological progression seen across the PIN-to-PCa transition, particularly between 6 and 8 months, we searched for cell-intrinsic changes in *TG-MYC^+^* epithelial cells that emerged over time. *TG-MYC^+^* cells separated into 6 distinct Leiden clusters, as visualized on UMAP (**Fig 1G**). Surprisingly however, these clusters did not correspond to the different timepoints across the PIN-to-PCa transition (**Fig 1H**). To more sensitively test for gene expression programs that might separate *TG-MYC^+^* cells by timepoint, we performed single-cell Hierarchical Poisson Factorization (scHPF), which identifies latent gene expression programs and assigns each cell a continuous loading for each program, allowing detection of subtle transcriptional signals. Consistent with the UMAP analysis, we failed to identify any transcriptional program that distinguished different timepoints across the PIN-to-PCa transition (**Fig 1I**). This lack of signal even under scHPF points to a striking degree of tumor cell-intrinsic transcriptional uniformity across timepoints. In summary, these findings suggest that the oncogenic *TG-MYC* transcriptional program in late-stage invasive PCa cells is already established at the preinvasive PIN stage and undergoes minimal remodeling during progression to invasive disease. In contrast, disease progression in other PCa models such as PtRP (*Pten^−/−^Rb^−/−^Trp53^−/−^*) is associated with large changes in tumor cell-intrinsic gene expression program upon cancer initiation.

### PIN-to-PCa transition is driven by macrophage infiltration

Puzzled by the lack of obvious tumor cell-intrinsic transcriptional programs specifically linked to invasion in PCa, we focused on cancer cell-extrinsic factors, noting that the initial UMAPs of whole prostate across all timepoints revealed an increase in immune cells in Hi-Myc tumors compared to wild-type prostates (particularly myeloid and T cells) (**Fig 1B-C, S1C**).

Focusing on the timepoint at which invasion begins (6 months), we observed that ∼60% of non-epithelial TME components are immune cells, of which ∼40% are myeloid cells with a predominance of macrophages based on expression of *Csf1r*, *Cd68*, *Adgre1*, *Aif1* (by scRNA-seq) and F4/80^+^ (by flow cytometry) (**Fig 2A-D, S2A-B**). Tumor-infiltrating F4/80^+^ macrophages were detectable in Hi-Myc mice as early as age 6 weeks, progressively increasing at 3, 6 and 8 months, eventually accounting for >40% of all CD45^+^ cells (**Fig 2E-F**).

**Figure 2.**
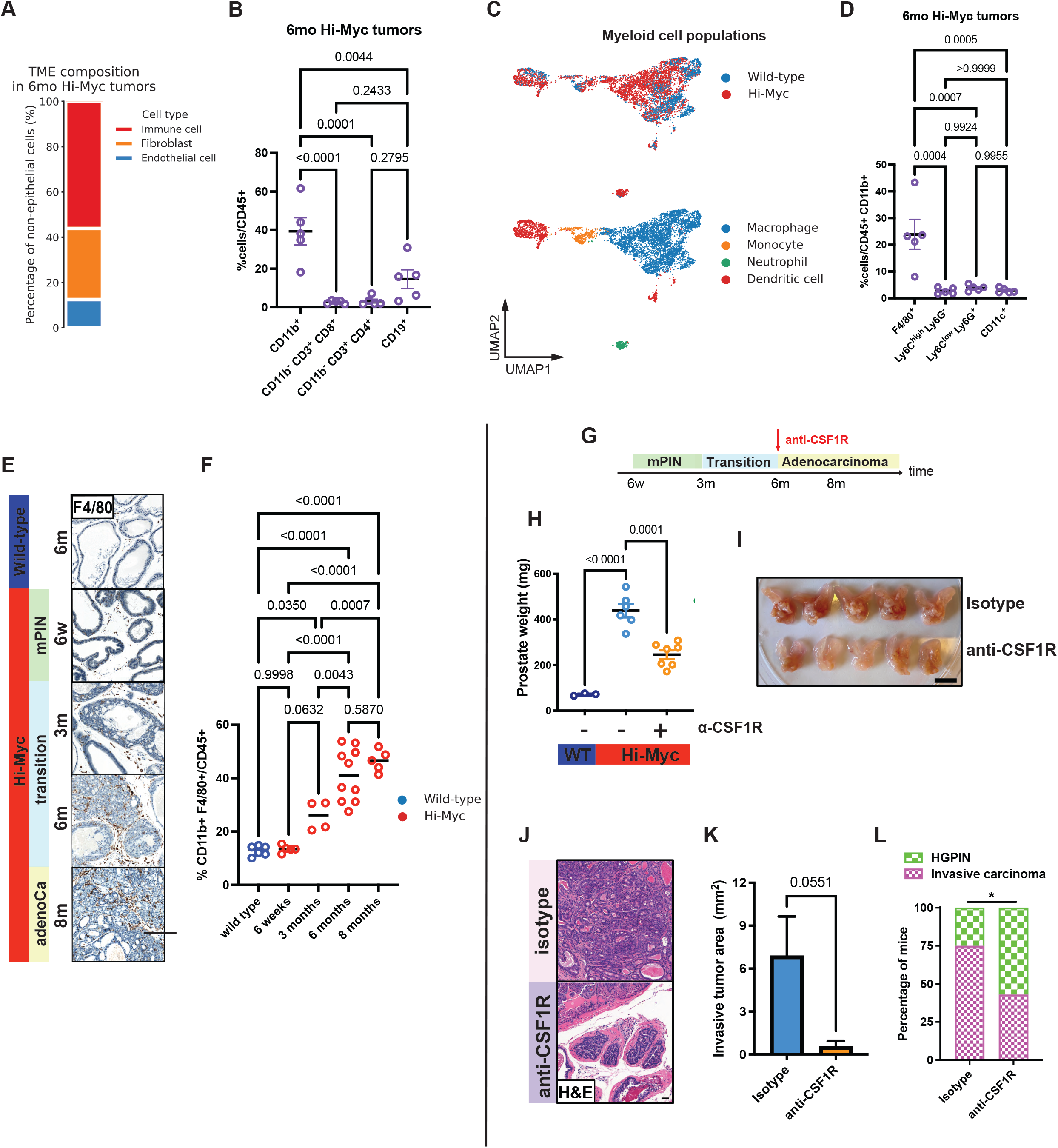
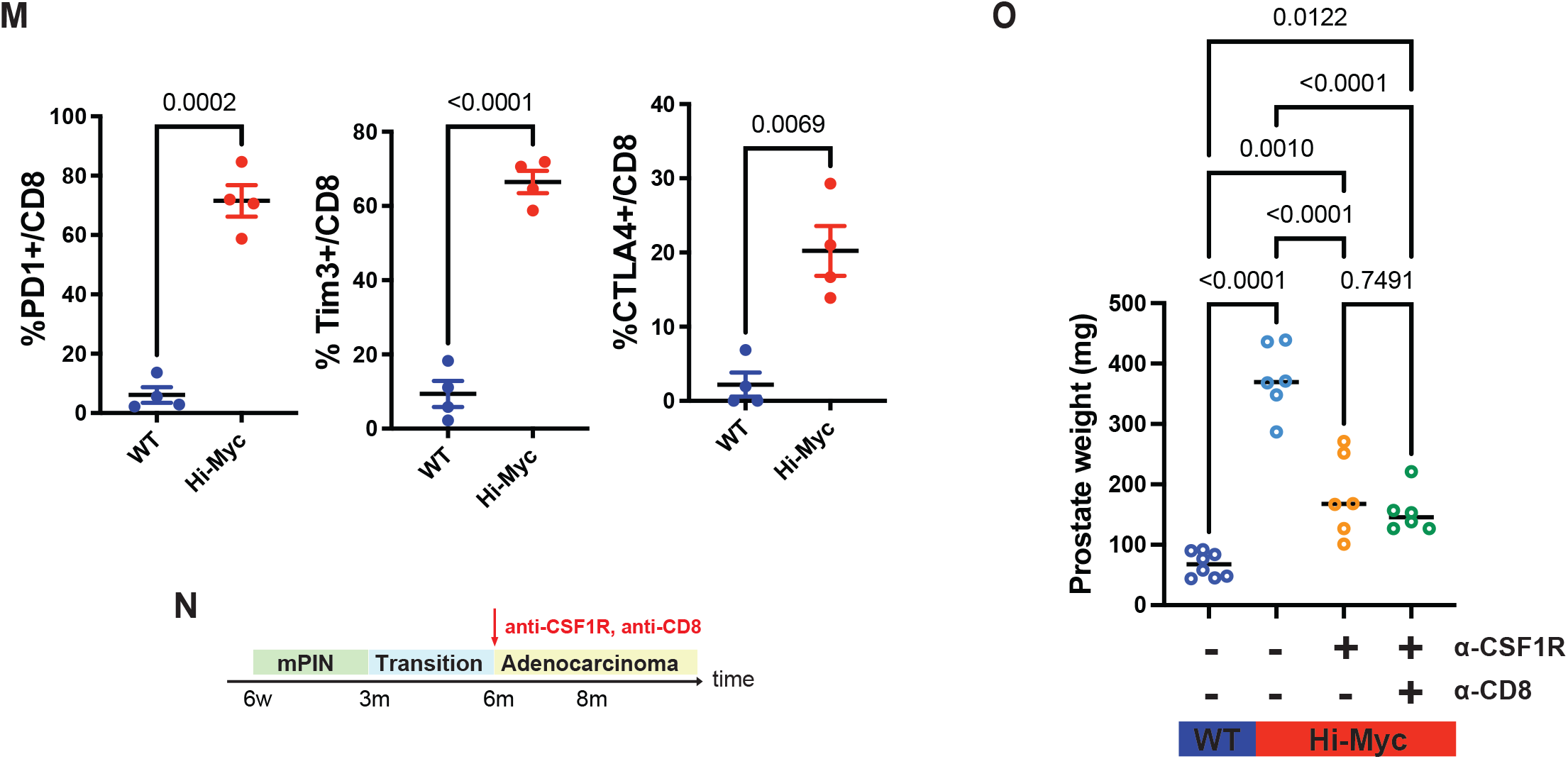
Macrophages drive the progression from PIN to invasive PCa. (A) Bar graph showing the proportion of non-epithelial cell populations in Hi-Myc prostates at early (6 weeks and 3 months) or late (6 and 8 months) timepoints. (B) Quantification of immune cell subsets in 6-month-old Hi-Myc tumors as determined by flow cytometry. One-way ANOVA; data are represented as mean ± s.d. (C) UMAP visualization of genotypes (top) and cell type annotations (bottom) within myeloid populations. (D) Quantification of myeloid cell subsets in 6-month-old Hi-Myc tumors as determined by flow cytometry. One-way ANOVA; data are represented as mean ± s.d. (E) Representative IHC staining for the macrophage marker (F4/80). Scale bar: 200 µm. (F) Quantification of macrophages by flow cytometry in wild-type and Hi-Myc tumors. One-way ANOVA; horizontal lines represent the mean. (G) Schematic of a 10-week anti-CSF1R antibody treatment regimen in Hi-Myc mice, initiated at 6 months of age. 6w, 6-week-old; 3m, 3-month-old; 6m, 6-month-old; 8m, 8-month-old. (H and I) Prostate weights (H) and representative images (I) following antibody treatment as outlined in (G). Scale bar: 10 mm. One-way ANOVA; data are represented as mean ± s.d. (J) Representative H&E images following antibody treatment. Scale bar: 50 µm. (K and L) Quantification of invasive area in tumors (K) and pathological tumor stage assessment (L) performed by an external clinical pathologist following anti-CSF1R antibody treatment of 6-month-old Hi-Myc mice. Unpaired two-tailed t-test; data are represented as mean ± s.d. (M) Quantification of T cells expressing exhaustion markers in prostates from 6-month-old wild-type (blue) and Hi-Myc (red) mice. Unpaired two-tailed t-test; data are represented as mean ± s.d. (N) Schematic of a 10-week anti-CSF1R ± anti-CD8 antibody treatment regimen in Hi-Myc mice, initiated at 6 months of age. 6w, 6-week-old; 3m, 3-month-old; 6m, 6-month-old; 8m, 8-month-old. (O) Prostate weights following antibody treatment of 6-month-old Hi-Myc mice as outlined in (N). One-way ANOVA; horizontal lines represent the mean.

This progressive increase in macrophage infiltration, particularly at the 6-month timepoint when PIN begins to transition to invasive PCa, raises the question of whether macrophages play a role in promoting invasion or accumulate as a response to tissue injury caused by invasion. To distinguish between these possibilities, we performed immunodepletion experiments using an antibody targeting CSF1R which is broadly expressed across tumor-infiltrating macrophages in Hi-Myc prostates (**Fig S2A**), beginning at 6 months and continuing through 8 months (**Fig 2G**). αCSF1R antibody treatment resulted in ∼60 percent depletion of F4/80^+^ intraprostatic macrophages and a ∼2-fold reduction in prostate weight, as well as a striking absence of visible tumor nodules in resected prostate lobes (**Fig 2H-I, S2C**). Histologically, αCSF1R antibody-treated mice prostates showed a ∼10-fold reduction in surface area showing tumor invasion, and more than half of the mice had no detectable evidence of invasion at 8 months (**Fig 2J-L**). To determine if macrophage depletion also impacts disease progression when invasive PCa is already present, we treated Hi-Myc mice with anti-CSF1R antibody at age 8 months (**Fig S2D**). As with the 6-month timepoint, we observed a significant decrease in prostate weight but no longer saw an effect on invasion (**Fig S2E-H**). Together, these data establish a pro-tumorigenic role of macrophages in the Hi-Myc model, particularly during the early stages of the PIN-to-PCa transition.

Abundant previous evidence has implicated myeloid cells in disease progression in PCa, particularly neutrophils in the context of *Pten* loss.^13,63,64^ Macrophages have been similarly implicated in lung, pancreatic, and breast cancer progression.^65–70^ Mechanistically, the myeloid infiltrates in these models are often associated with impaired CD8^+^ T cell function, hence the term myeloid-derived suppressor cells, which can interfere with tumor immune surveillance.

Along these lines, we observed increased numbers of T cells in Hi-Myc tumors, with roughly equal proportions of CD4^+^ and CD8^+^ cells (**Fig 1B-C, 2B**). Furthermore, a large fraction of CD8^+^ T cells co-expressed exhaustion markers such as PD1 (∼70%), TIM3 (∼60%), and CTLA4 (∼20%), and expression of these markers declined 2 to 3-fold following one week of macrophage depletion (**Fig 2M, S2I**). Having established that macrophages in the Hi-Myc model also generate an immunosuppressive TME, we asked whether the protective effect of macrophage depletion (reduced invasion) is immune-mediated by co-depleting CD8^+^ T cells and macrophages. Surprisingly, depleting CD8^+^ T cells did not reverse the protective effect of macrophage depletion on tumor burden (**Fig 2N-O, S2J**), suggesting that the role of macrophages in promoting the PIN-to-PCa transition is independent of CD8^+^ T cells.

### IL-1β is upregulated in intraprostatic macrophages and required for invasion

To gain insight into how the macrophages that infiltrate the prostates of Hi-Myc mice might promote invasion independent of CD8^+^ T cells, we used our transcriptomic data to further define the macrophage subtypes present in Hi-Myc mice. We noted abundant expression of *Trem2* (**Fig 3A**), consistent with data from other epithelial cancer models.^71,72^ TREM2 senses cell debris involved in lipid sensing and regulation of phagocytosis but is also linked with immune suppression in cancer.^72–74^ In contrast, macrophages expressing markers of tissue-resident macrophages such as *Folr2*, *Lyve1* and *Timd4*^75–77^ were limited to minor subsets. Interestingly, *Spp1* expression was more limited, in contrast to advanced human prostate cancers in which tumor-infiltrating macrophages are characterized by high co-expression of TREM2 and SPP1,^38,78^ perhaps indicative of distinct macrophage biology in early-stage disease.

**Figure 3.**
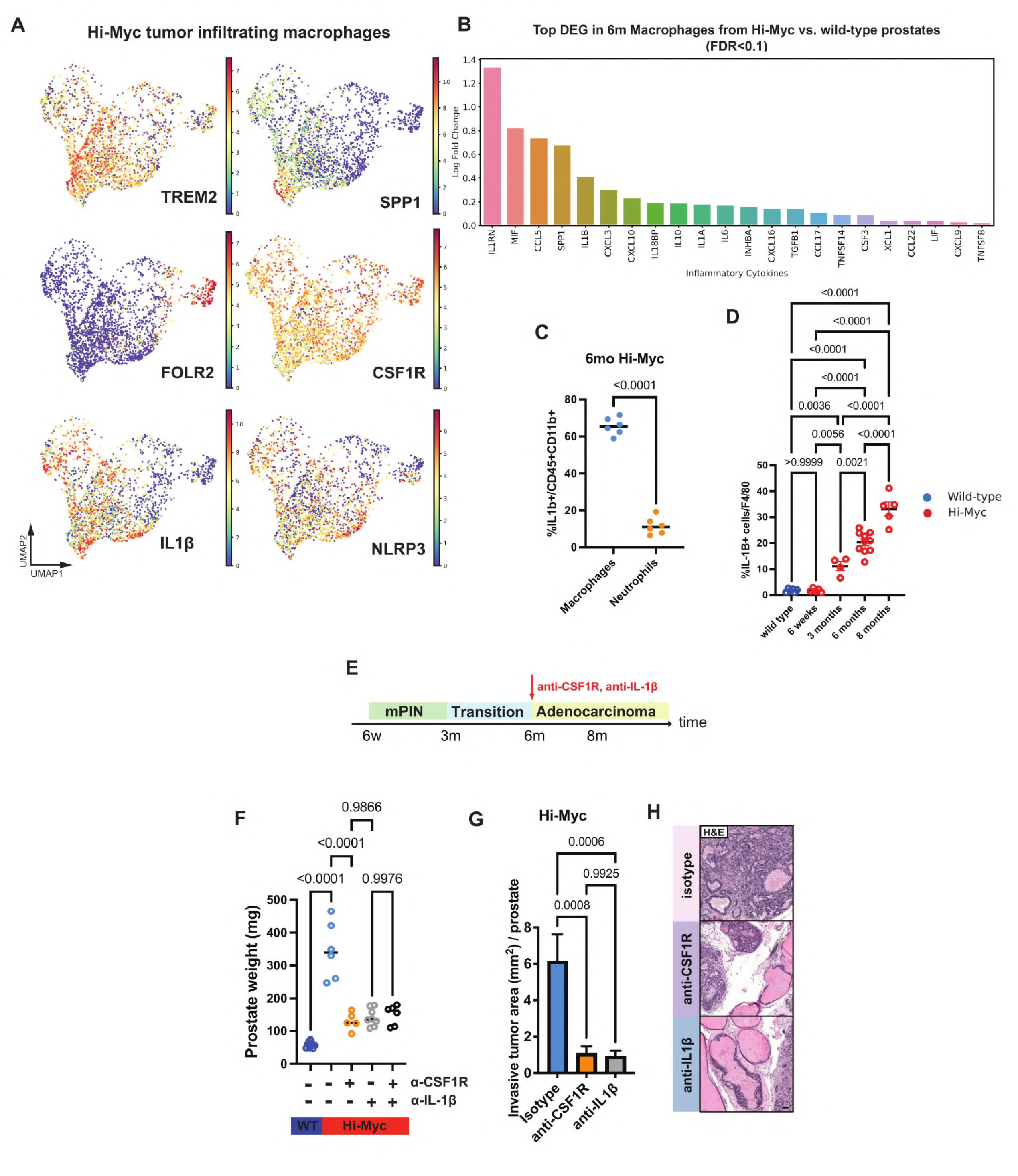
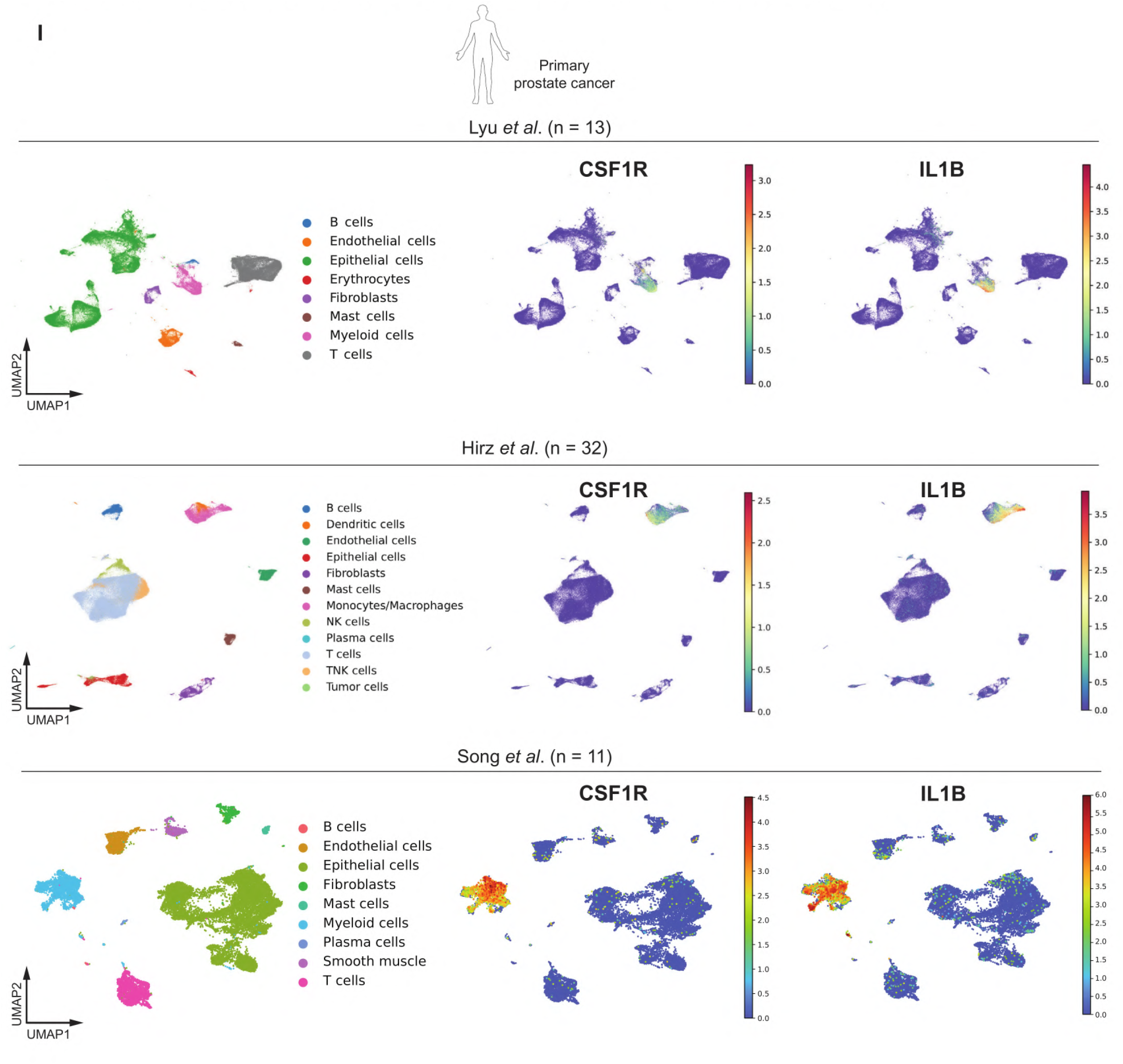
Macrophages accelerate prostate cancer development via IL-1β signaling. (A) UMAP visualization of expression of indicated genes in Hi-Myc tumor infiltrating macrophages. (B) Differentially expressed inflammatory cytokines in macrophages from 6-month-old Hi-Myc prostates compared with those from age-matched wild-type prostates (FDR<0.1). (C) Quantification of IL-1β^+^ macrophages (CD45^+^ CD11b^+^ Ly6C^low^ Ly6G^-^ F4/80^+^) and IL-1β^+^ neutrophils (CD45^+^ CD11b^+^ Ly6C^-^ Ly6G^+^) in 6-month-old Hi-Myc prostates. Unpaired two-tailed t-test; horizontal lines represent the mean. (D) Quantification of IL-1β^+^ macrophages (CD45^+^ CD11b^+^ Ly6C^low^ Ly6G^-^ F4/80^+^) in wild-type and Hi-Myc prostates across indicated ages. One-way ANOVA; data are represented as mean ± s.d. (E) Schematic of a 10-week treatment regimen with anti-CSF1R or anti-IL-1β antibodies in Hi-Myc mice, initiated at 6 months of age. 6w, 6-week-old; 3m, 3-month-old; 6m, 6-month-old; 8m, 8-month-old. (F) Prostate weights following antibody treatment of 6-month-old Hi-Myc mice. One-way ANOVA; horizontal lines represent the mean. (G) Quantification of invasive tumor area following antibody treatment of 6-month-old Hi-Myc mice. One-way ANOVA; data are represented as mean ± s.d. (H) Representative H&E images following antibody treatment. Scale bar: 50 µm. (I) UMAP visualization of *CSF1R* and *IL1B* expression across cell types in primary human prostate cancer scRNA-seq datasets.

To explore how these TREM2^+^ macrophages promote PCa invasion independent of CD8^+^ T cells, we examined differentially expressed genes in macrophages from Hi-Myc versus wild-type prostates and noted striking upregulation of *Il1rn* and *Il1b* as well as other inflammatory cytokines such as *Mif* and *Ccl5* (**Fig 3B**). Because IL-1β has been previously linked with tumor progression in lung and pancreatic cancer,^35,65,79^ we directly examined IL-1β protein expression by flow cytometry. Notably, expression was exclusive to macrophages (and not neutrophils) (**Fig 3C**) and strikingly upregulated in prostates of Hi-Myc mice, with a progressive increase in percentage of IL-1β^+^ macrophages during the PIN-to-PCa transition (**Fig 3D**). To determine whether IL-1β plays a role in tumor progression, we treated 6-month-old Hi-Myc mice with an IL-1β-neutralizing antibody for 10 weeks, following the schedule used earlier for macrophage depletion with the anti-CSF1R antibody (**Fig 3E**) and observed a comparable reduction in tumor weight (∼3-fold) and decrease in tumor invasion (∼6-fold) (**Fig 3F-H**). Combination treatment of anti-IL-1β and anti-CSF1R antibodies resulted in a reduction in prostate weight to a degree comparable to that seen with either single agent (**Fig 3F**), supporting the hypothesis that IL-1β release from tumor-infiltrating macrophages is a critical effector of the pro-invasion phenotype.

To determine if these findings in Hi-Myc mice extend to prostate cancers initiated by other oncogenic events, we analyzed scRNA-seq data from two models in which tumors are initiated by loss of tumor suppressor genes commonly mutated in human prostate cancer (*Pten^-/-^; Rb1^-/-^* and *Pten^-/-^; Rb1^-/-^; Trp53^-/-^*). In both cases, *Il1b* and *Il1rn* were among the top upregulated genes in tumor-infiltrating macrophages (**Fig S3A-B**), suggesting the role of IL-1β in prostate cancer progression is not genotype-specific. To extend the analysis to human prostate cancer, we analyzed scRNA-seq data from human prostate cancer prostatectomy specimens (56 primary samples across 3 independent cohorts)^38–40^ and found clear evidence of *IL1B* expression in *CSF1R*^+^ macrophages in tumors from all three cohorts but not in normal human prostate tissue (**Fig 3I, S3C**). Taken together, the evidence across several mouse prostate cancer models and human prostate cancer datasets implicates IL-1β production by tumor-infiltrating macrophages as a driver of early prostate cancer development.

### IL-1β acts through the tumor microenvironment to promote prostate cancer progression

Having demonstrated a critical role for macrophage-derived IL-1β in PCa invasion, we next sought to define the cellular target of IL-1β action. Prior work in pancreatic cancer defined a signaling loop whereby IL-1β produced by macrophages acts directly on tumor cells.^65^ Similarly, in lung cancer, IL-1β from macrophages has been shown to enhance the regenerative capacity of *Kras*-mutant AT2 cells through direct engagement of IL-1 receptor (IL-1R1).^35,80^ To determine if IL-1β also acts directly on tumor cells in prostate cancer, we developed an organoid transplantation model in which primary prostate organoids were engineered to express high levels of a stabilized form of MYC (*Myc^T58A^*, mimicking the Hi-Myc GEMM) together with *Trp53* deletion to enhance engraftment (hereafter called PM), then transplanted orthotopically into syngeneic hosts (see Methods) (**Fig 4A**). After a single *in vivo* passage, PM orthografts consistently developed locally invasive adenocarcinomas (within 3 weeks) accompanied by infiltration by IL-1β^+^ macrophages (**Fig 4B-C, S4A-B**), mirroring the pre- and post-invasion phenotype in Hi-Myc mice but now with a timescale and platform amenable to higher throughput genetic perturbation. Importantly, tumor progression in the orthotopic PM model is also IL-1 signaling-dependent as revealed by treatment with an IL-1R1-blocking antibody (**Fig 4D-E**).

**Figure 4.**
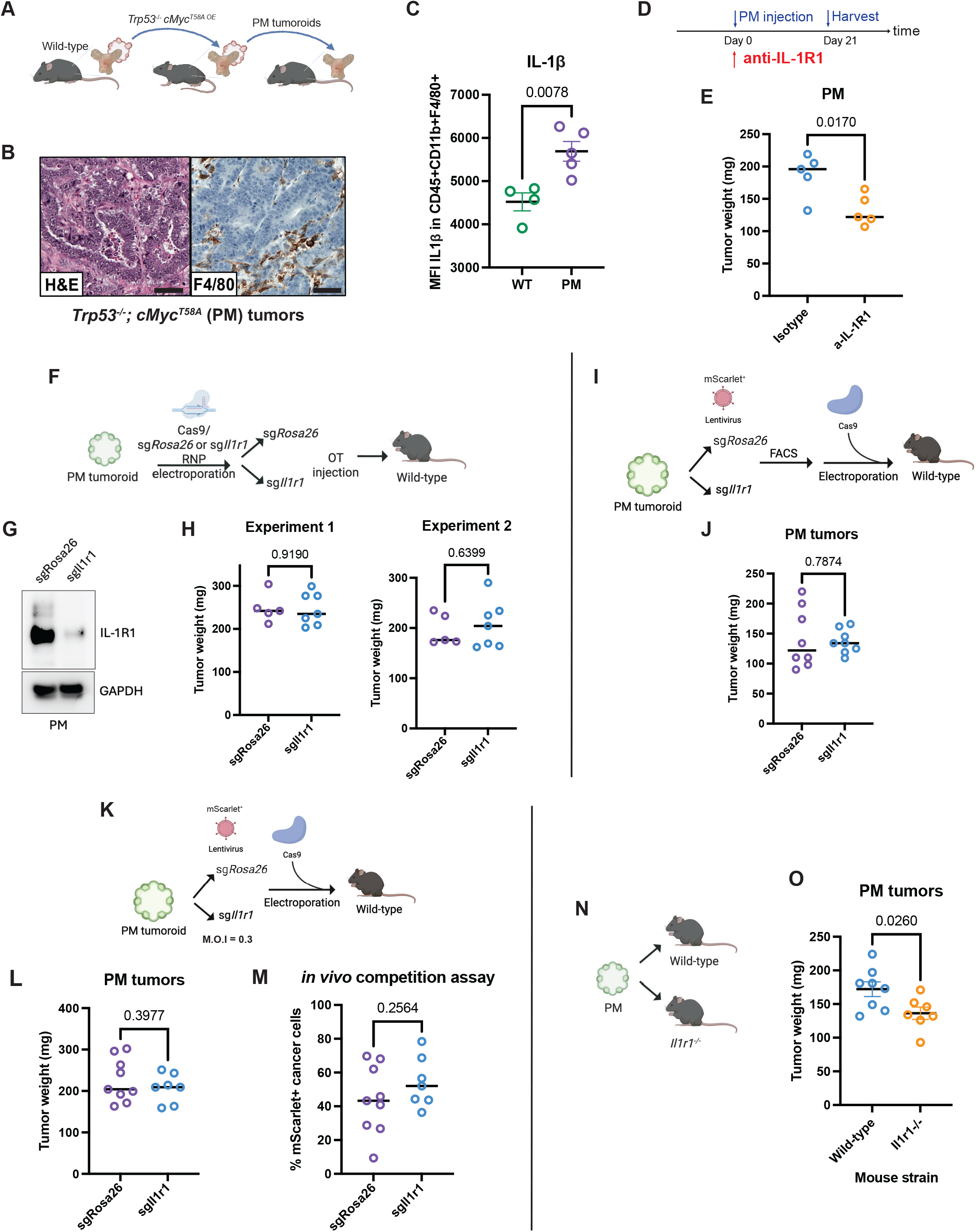
IL-1β promotes prostate cancer progression through a cancer cell-extrinsic mechanism. (A) Schematic outlining the generation of PM tumoroids from mouse prostate organoids. (B) Representative H&E and F4/80 immunostaining of orthotopic PM tumors. Scale bar: 50 µm. (C) Mean fluorescence intensity of IL-1β in macrophages from wild-type prostates (green) and macrophages in PM tumors (purple). Unpaired two-tailed t-test; data are represented as mean ± s.d. (D) Schematic outlining PM tumoroid injection and antibody treatment regimen. (E) PM tumor weights following antibody treatment as described in H. Unpaired two-tailed t-test; horizontal lines represent the mean. (F) Schematic illustrating the electroporation introduction of Cas9-sgRNA RNP into PM tumoroids for *in vivo* knockdown studies. (G) Western blot confirming *Il1r1* knockdown efficiency shown in (F). GAPDH was used as a loading control. (H) PM tumor weights from (F) in two independent, biological replicate experiments (Experiment 1, 2). Unpaired two-tailed t-test; horizontal lines represent the mean. (I) Schematic illustrating the introduction of lentiviral sgRNA and Cas9 by electroporation into PM tumoroids for *in vivo* knockdown studies. (J) PM tumor weights from (I). Unpaired two-tailed t-test; horizontal lines represent the mean. (K) Schematic illustrating the introduction of Cas9-sgRNA system into PM tumoroids for *in vivo* competition assays. (L) PM tumor weights from (K). Unpaired two-tailed t-test; horizontal lines represent the mean. (M) Percent change of mScarlet^+^ cancer cells in post-graft tumors. Unpaired two-tailed t-test; horizontal lines represent the mean. (N) Schematic illustrating the introduction of PM tumoroids into wild-type or *Il1r1^-/-^* mice. (O) PM tumor weights from (N). Unpaired two-tailed t-test; data are represented as mean ± s.d.

To determine if IL-1β-driven tumor progression is mediated by direct action on tumor cells, we deleted *Il1r1* in PM tumoroids by electroporation of CRISPR-Cas9 ribonucleoprotein (RNP), confirmed loss of IL-1R1 protein expression by western blot, and measured tumor size 3 weeks after orthotopic transplantation (**Fig 4F-G**). Surprisingly, *Il1r1* deletion had no effect on tumor engraftment or weight in two independent, biological replicate experiments (**Fig 4H**). To confirm this result, we repeated the experiment but now introduced sg*Il1r1* by lentivirus with a cis-linked mScarlet fluorescence marker to enable enrichment of sg*Il1r1-*infected cells by FACS. Again, we observed no difference in tumor weight following *Il1r1* knockdown (**Fig 4I-J**). As a final test, we performed an *in vivo* competition experiment in which PM tumoroids were infected with mScarlet^+^ lentivirus (sg*Il1r1* or control sg*Rosa26*) at low multiplicity of infection (MOI). We then scored the percentage of mScarlet^+^ cancer cells in the tumors that developed following orthotopic transplantation, reasoning that sg*Il1r1*-infected cells should dropout if IL-1β promotes tumor progression through direct engagement of IL-1R1 on tumor cells. However, consistent with the earlier approaches, we observed no loss of mScarlet positivity following *Il1r1* knockdown (**Fig 4K-M**), confirming that IL-1β does not promote tumor progression through direct action on cancer cells. To formally test the alternative hypothesis that IL-1β impacts tumor growth indirectly, we repeated the PM tumoroid transplantation experiment using syngeneic *Il1r1^-/-^* (germline deleted) mice as hosts. Indeed, tumor growth in *Il1r1^-/-^* mice was impaired relative to wild-type mice (**Fig 4N-O**), consistent with the results using the IL-1R1 blocking antibody (**Fig 4E**), now providing clear evidence that IL-1β promotes tumor progression through an indirect mechanism.

### Tumor progression is driven by an IL-1β (macrophage) -> IL-6 (fibroblast) -> tumor cell circuit

To identify the host cell responsible for IL-1β-dependent tumor progression, we examined *Il1r1* mRNA levels across all cell types within the TME of Hi-Myc tumors and noted the highest expression in fibroblasts (**Fig 5A**). This finding was confirmed at the protein level with robust IL-1R1 expression in fibroblasts isolated from 6-month-old Hi-Myc tumors, at levels higher than in prostate fibroblasts from age-matched wild-type mice (**Fig S5A**). To search for evidence of IL-1R1 activation in Hi-Myc fibroblasts, we compared their gene expression with that of fibroblasts from wild-type prostates and noted upregulation of several myeloid chemokines (*Ccl2*, *Ccl7*, *Cxcl1*, *Cxcl2*) and *Il6,* a known growth factor for many epithelial tumor types including prostate,^81^ as well as enrichment of inflammatory signaling pathways by GSEA (**Fig 5B, S5B**). Again, we confirmed these findings at the protein level in the conditioned media of fibroblasts from Hi-Myc or wild-type prostates propagated *ex vivo* (**Fig 5C-D**).

**Figure 5.**
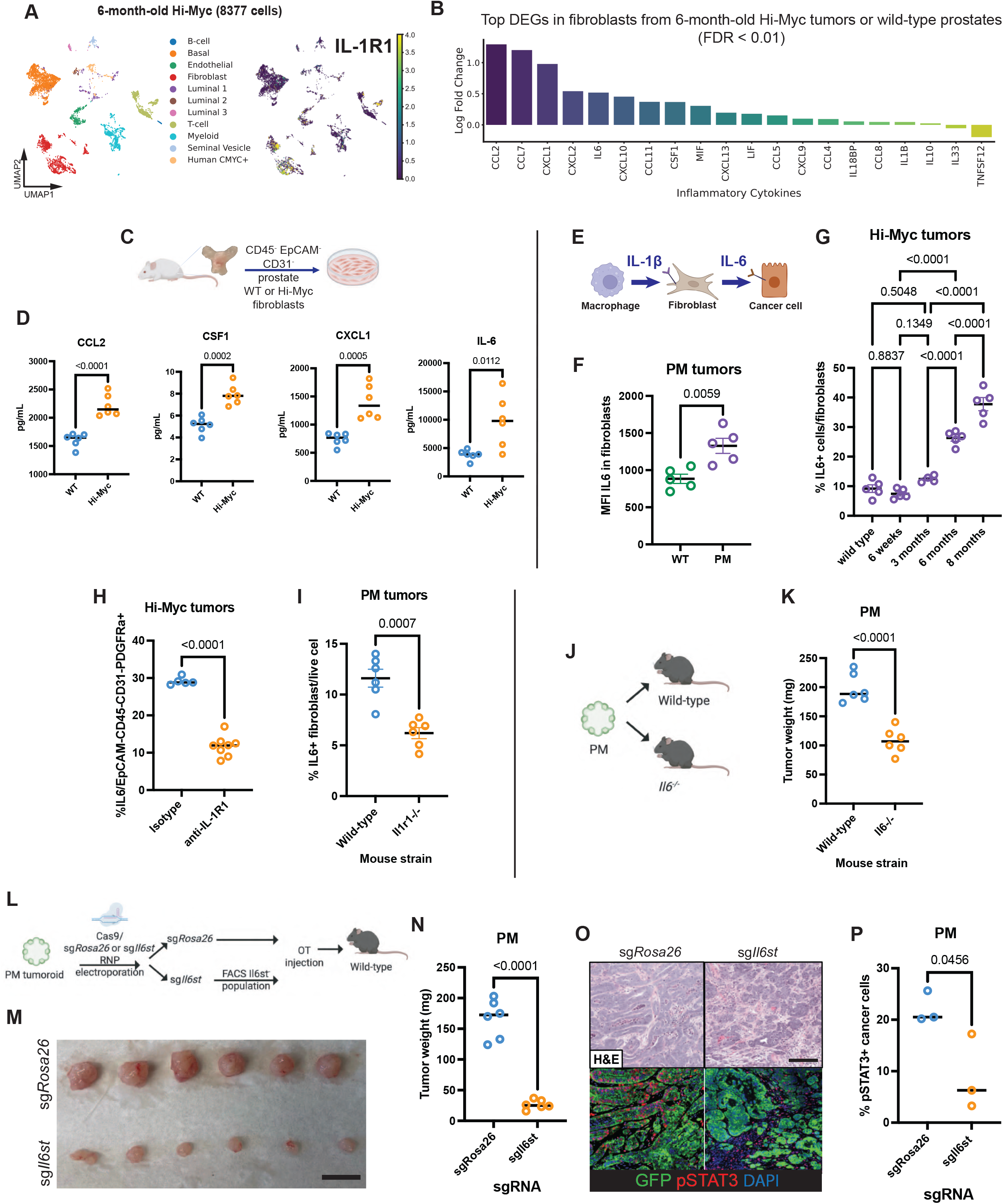
Fibroblast-derived IL-6, induced by IL-1β signaling, promotes tumor progression. (A) UMAP visualization of cell types (left) and *Il1r1* gene expression (right) in 6-month-old Hi-Myc prostates (n=8,377). (B) Differentially expressed inflammatory cytokines in fibroblasts from 6-month-old Hi-Myc prostates compared with those from wild-type prostates (FDR<0.01). (C) Schematic of the isolation workflow for prostate fibroblasts from wild-type or Hi-Myc prostates for *ex vivo* studies. (D) Protein levels in conditioned media following 24-hr incubation of wild-type or Hi-Myc prostate fibroblasts. Unpaired two-tailed t-test; horizontal lines represent the mean. (E) Schematic summarizing the IL-1β (macrophage) -> IL-6 (fibroblast) -> tumor cell signaling circuit. (F) Mean fluorescence intensity of IL-6 expression in fibroblasts from wild-type prostates (green) and fibroblasts in PM tumors (purple). Unpaired two-tailed t-test; data are represented as mean ± s.d. (G) Quantification of IL-6^+^ cells in fibroblasts (CD45^-^ EpCAM^-^ CD31^-^ PDGFRα^+^) in wild-type and Hi-Myc prostates across different mouse ages. One-way ANOVA; data are represented as mean ± s.d. (H) Quantification of IL-6^+^ cells in Hi-Myc fibroblasts (CD45^-^ EpCAM^-^ CD31^-^ PDGFRα^+^) following antibody treatment. Unpaired two-tailed t-test; data are represented as mean ± s.d. (I) Quantification of IL-6^+^ cells in PM fibroblasts (CD45^-^ EpCAM^-^ CD31^-^ PDGFRα^+^) after PM tumoroid injection into wild-type or *Il1r1^-/-^* mice. Unpaired two-tailed t-test; data are represented as mean ± s.d. (J) Schematic illustrating the introduction of PM tumoroids into wild-type or *Il6^-/-^* mice.| (K) PM tumor weights from (J). Unpaired two-tailed t-test; horizontal lines represent the mean. (L) Schematic illustrating the electroporation introduction of Cas9-sgRNA RNP into PM tumoroids for *in vivo* knockdown studies. (M and N) Representative images (M) and tumor weight quantification (N) of PM tumors from the experiment outlined in (L). Scale bar: 10 mm. Unpaired two-tailed t-test; horizontal lines represent the mean. (O and P) Representative H&E and multiplex immunofluorescence images (O) and quantification of phospho-STAT3^+^ cancer cells (GFP^+^) (P). Scale bar: 100 µm. Unpaired two-tailed t-test; horizontal lines represent the mean.

Among the many chemokines/cytokines upregulated in fibroblasts from Hi-Myc mice, the fact that IL-6 is a direct transcriptional target of IL-1 signaling^82^ and has a known role in directly promoting epithelial tumor growth^81^ nominates it as a compelling candidate to explain how IL-1β promotes tumor progression indirectly (e.g., macrophage -> IL-1β -> fibroblast-> IL-6 -> tumor) (**Fig 5E**). Consistent with this hypothesis, expression of IL-6 protein in fibroblasts from Hi-Myc prostates, as well as from PM orthografts, progressively increased over time (**Fig 5F-G**), paralleling the increase in IL-1β protein expression seen in macrophages at precisely the same timepoints (**Fig 3D**). Although correlative, these data support the existence of an intraprostatic IL-1β (macrophage) -> IL-6 (fibroblast) circuit that subsequently drives tumor progression. As a first test of this hypothesis, we repeated the measurements of IL-6 protein expression in prostatic fibroblasts but now in the setting of IL-1R1 inhibition, reasoning that interruption of the circuit at the level of IL-1R1 should impair downstream expression of IL-6 by fibroblasts. Indeed, this was exactly the case in Hi-Myc mice treated with an IL-1R1-blocking antibody and in PM orthografts engrafted into *Il1r1^-/-^* host mice (**Fig 5H-I**). Having confirmed the existence of an IL-1β -> IL-6 circuit, we next asked if downstream interruption of the circuit at the level of IL-6 impairs tumorigenicity. Remarkably, PM tumoroids orthotopically transplanted into syngeneic *Il6^-/-^* hosts were significantly smaller than those transplanted into wild-type hosts (**Fig 5J-K**), mirroring the results seen through upstream interruption of the circuit by transplantation into *Il1r1^-/-^* mice (**Fig 4N-O**). Because IL-6 can be expressed by immune cells, tumor cells as well as fibroblasts,^83,84^ we examined IL-6 expression across all these cell types in Hi-Myc prostates and in PM orthografts and identified fibroblasts as the main source of IL-6 (**Fig S5C-D**), providing further validation of the IL-1β (macrophage) -> IL-6 (fibroblast) circuit model.

To determine the target of IL-6 signaling in our prostate models, we surveyed all cell types in the TME of Hi-Myc prostates and noted broad expression of both subunits of the IL-6 receptor complex (*Il6ra*, *Il6st*) (**Fig S5E-F**), consistent with known biology of IL-6 signaling across multiple cell types. To address the question of which of these cell types is critical for tumor progression in our prostate models, we first performed immunohistochemical staining for the IL-6 activation marker phospho-STAT3 and noted robust staining of tumor cells in the PM orthograft model (**Fig S5G**), suggesting tumor cells may be the key IL-6 target cells. To test this hypothesis directly, we deleted *Il6st* in PM tumoroids using CRISPR, followed by orthotopic injection into wild-type mice (**Fig 5L**). Remarkably, *Il6st* deletion resulted in a ∼6-fold reduction in tumor burden as well as a profound decrease in downstream phospho-STAT3 level (**Fig 5M-P**). Taken together, these experiments provide strong evidence for a TME circuit in which IL-1β secretion by macrophages induces IL-6 expression in prostate fibroblasts, which promotes prostate cancer progression through direct activation of IL-6R signaling in tumor cells.

### Evidence for an IL-1β (macrophage) -> IL-6 (fibroblast) circuit in other (non-MYC-driven) prostate models and in human prostate and lung adenocarcinoma

To determine if the IL-1β-IL-6 circuit present in Hi-Myc mice is also seen in prostate cancer models driven by alternative genomic alterations, we turned to the PtR and PtRP models discussed earlier, each of which showed elevated levels of IL-1β in tumor-infiltrating macrophages (**Fig S3A-B**). As with Hi-Myc mice, fibroblasts in the prostates of PtR and PtRP mice expressed elevated levels of *Il6* as well as the myeloid chemokines *Ccl2, Ccl7, Cxcl1* and *Cxcl2* (**Fig S6A-B**), evidence that the macrophage -> IL-1β -> fibroblast-> IL-6 circuit is not unique to a specific genotype.

To extend the analysis to human prostate cancer, we used scRNA-sequencing gene signatures derived from tumor-infiltrating mouse macrophages (reflective of IL-1β activation) and from tumor-associated mouse prostate fibroblasts (reflective of upregulated IL-6 and myeloid chemokine expression) to interrogate primary human prostate cancer datasets. We stratified patients into two groups based on clinical prognosis: a lower-risk group (Gleason score 3+4 or lower) and a higher-risk group (Gleason score 4+3 or higher).^85^ Gleason score is broadly used to assess tumor aggressiveness in patients. Higher gene signature scores for both Hi-Myc macrophages and Hi-Myc fibroblasts were significantly associated with higher-risk Gleason group (**Fig 6A**). Importantly, the two signature scores were positively and significantly correlated with each other not only in human prostate cancer but also in lung adenocarcinoma (**Fig 6B-C**), providing evidence that the IL-1β -> IL-6 circuit revealed in mouse prostate cancer models is present in patients. The lung cancer result is particularly striking based on the strength of the association (r=0.619) as well as clinical evidence linking IL-1β-neutralizing antibody therapy with a reduction in incident lung cancer.^86^

**Figure 6.**
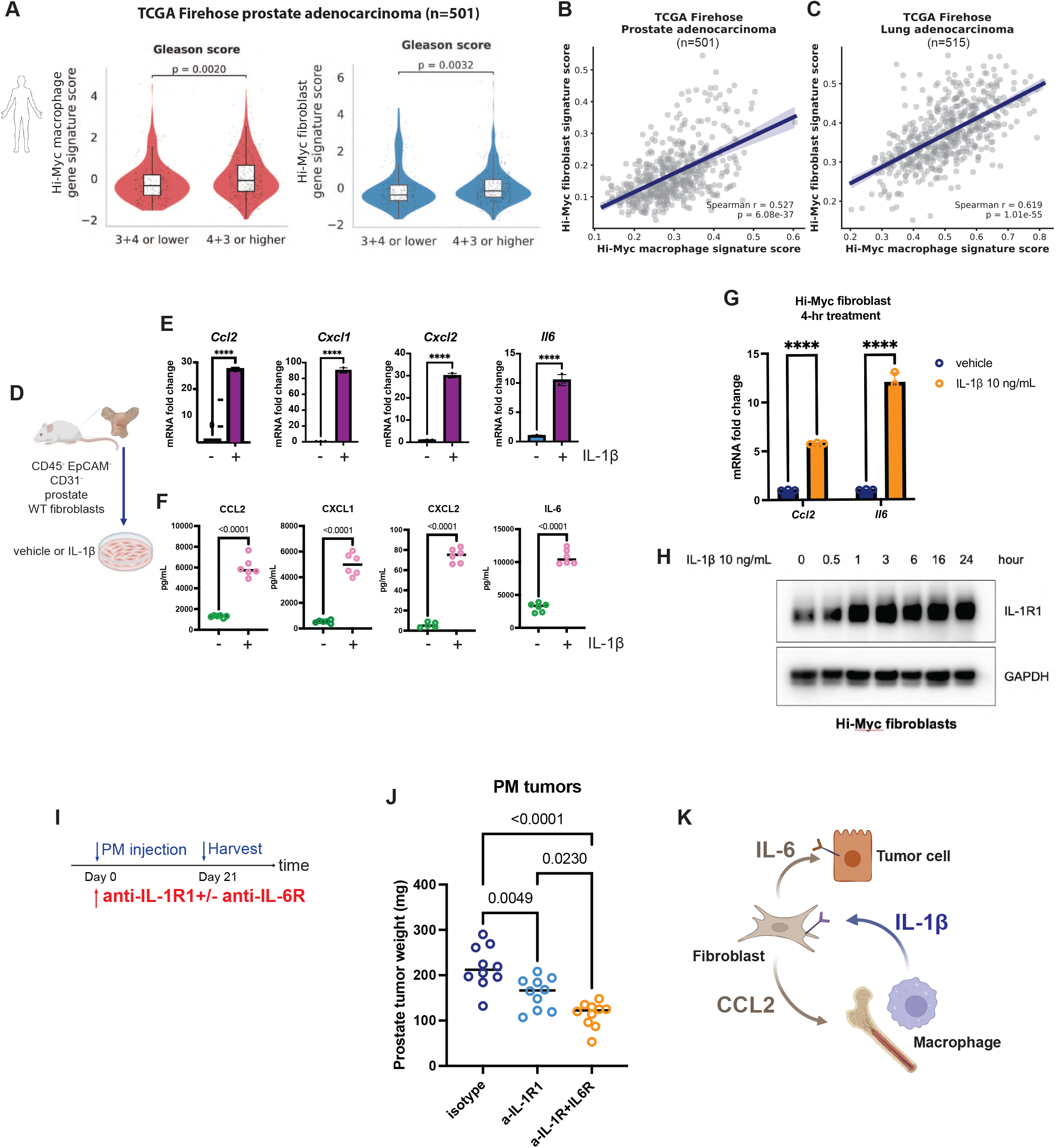
The IL-1β (macrophage) – IL-6 (fibroblast) circuit is conserved across prostate cancer models and human prostate and lung cancer. (A) Correlation between Gleason score and Hi-Myc macrophage (left) or fibroblast (right) gene signature score in primary prostate cancer patients from The Cancer Genome Atlas (TCGA; n=501). Unpaired two-tailed t-test; data are represented as mean ± s.d.| (B and C) Correlation between Hi-Myc macrophage and fibroblast gene signature scores in human primary prostate cancer (B) or lung cancer (C) (The Cancer Genome Atlas Firehose). (D) Schematic of the isolation workflow for prostate fibroblasts from wild-type prostates for *ex vivo* studies. (E) mRNA levels of target genes in wild-type prostate fibroblasts after 4-hr IL-1β treatment *ex vivo*. Unpaired two-tailed t-test; data are represented as mean ± s.d. (F) Protein levels of target genes in conditioned media following 24-hr IL-1β treatment of wild-type fibroblasts. Unpaired two-tailed t-test; data are represented as mean ± s.d. (G) mRNA levels of target genes in 6-month-old Hi-Myc prostate fibroblasts after 4-hr IL-1β treatment *ex vivo*. Unpaired two-tailed t-test; data are represented as mean ± s.d. (H) Western blot showing IL-1R1 levels in prostate fibroblasts from 6-month-old Hi-Myc mice following a time-course of IL-1β treatment. GAPDH was used as a loading control. (I) Schematic outlining PM tumoroid injection and antibody treatment regimen. (J) PM tumor weights from (I). Unpaired two-tailed t-test; horizontal lines represent the mean. (K) Schematic summarizing the pro-tumorigenic IL-1β/IL-6 signaling axis within the prostate tumor microenvironment.

Recognizing the broader relevance of the IL-1β -> IL-6 circuit, we next considered potential mechanisms by which the circuit gets established, reasoning that normal prostate fibroblasts may require reprogramming in response to IL-1β stimulation. However, we found that fibroblasts freshly isolated from wild-type mouse prostates responded robustly to recombinant IL-1β stimulation *ex vivo*. Specifically, we observed a robust increase in *Ccl2, Cxcl1, Cxcl2*, and *Il6* mRNA levels within 4 hours, as well as release of these cytokines into fibroblast-conditioned media at 24 hours (**Fig 6D-F**), indicating that normal prostate fibroblasts can respond to IL-1β in the absence of reprogramming. Interestingly, both *Ccl2* and *Il6* were further upregulated after IL-1β exposure in fibroblasts isolated from 6-month-old Hi-Myc mouse prostates, consistent with enhanced IL-1β responsiveness at the PIN stage (**Fig 6G**). IL-1β treatment also resulted in an increase in IL-1R1 protein expression within 24 hours (**Fig 6H**), consistent with a feedforward loop whereby IL-1β upregulates expression of its own receptor, as previously described in colorectal cancer,^87^ further amplifying the effects of IL-1β -> IL-6 circuit.

As discussed earlier, clinical results evaluating the therapeutic utility of IL-1β neutralization in patients with established lung cancer (the CANOPY trials with canakinumab) have been disappointing.^44,45,47^ With evidence of an IL-1β -> IL-6 circuit in prostate cancer (and potential relevance in lung cancer), we asked whether combined IL-1β + IL-6 inhibition might be more effective than IL-1β blockade alone in our prostate cancer model. Indeed, treatment with anti-IL-1R1 + IL-6R blockade resulted in a significant (nearly 2-fold) reduction in tumor volume in PM orthografts compared to IL-1R1 blockade alone (**Fig 6I-J**). Collectively, these results support further exploration of combined IL-1β and IL-6 neutralization to block a pro-tumorigenic TME signaling circuit across prostate and lung cancer models at early stages of disease (**Fig 6K**).

## Discussion

Through single-cell characterization of a widely studied mouse prostate cancer model (Hi-Myc) in which a prolonged preneoplastic (PIN) stage precedes development of invasive adenocarcinoma, we reveal that IL-1β^+^ tumor-infiltrating macrophages play a critical role in driving this progression. In contrast to earlier work showing that IL-1β from tumor-infiltrating macrophages stimulates cancer progression in lung and pancreatic cancer models by directly acting on tumor cells,^35,65^ in prostate cancer IL-1β activates a TME signaling circuit, acting initially on prostate fibroblasts to stimulate secretion of IL-6, which then drives cancer progression by activating the IL-6 receptor on tumor cells. Considering prior work documenting autocrine IL-6 production by prostate cancer cells in certain contexts,^81,88^ the fact that IL-6 from fibroblasts also drives prostate cancer progression is not surprising. However, our discovery of an IL-1β (macrophage) -> IL-6 (fibroblast) -> tumor circuit within the TME is novel. Furthermore, this circuit is likely relevant beyond the Hi-Myc model based on our transcriptomic analysis of other (non-MYC-driven) prostate cancer models as well as human prostate and lung adenocarcinoma datasets. Finally, we show that combined IL-1β + IL-6 blockade is more effective than IL-1β blockade alone, providing preclinical proof of concept that targeting a chemokine circuit at multiple levels may be more effective than single cytokine blockade.

Having demonstrated the existence of the IL-1β -> IL-6 TME circuit, an important next question is how it becomes activated during the early stages of cancer initiation. As a first step, we focused on the prostate fibroblasts that receive the IL-1β signal from macrophages. Despite abundant evidence that fibroblasts are reprogrammed during epithelial cancer initiation,^89^ we find that normal prostate fibroblasts are already primed to respond to IL-1β (**Fig 6D-F**). If fibroblast reprogramming is not required for circuit activation, what is? One possibility is that tissue-resident macrophages in the normal prostate secrete IL-1β after sensing a currently unknown signal from tumor cells produced early following oncogene activation. The subsequent action of IL-1β on adjacent fibroblasts then activates IL-6, as well as myeloid chemokines (CCL2, CCL7, CXCL1, CXCL2) that fuel recruitment of monocyte-derived macrophages to infiltrate the prostate, generating a feedforward loop that further increases the abundance of IL-1β and IL-6 in the TME. Alternatively, fibroblasts (not macrophages) may be first to sense an early signal of oncogene activation and then, through production of myeloid chemokines, recruit monocyte-derived macrophages to fuel subsequent IL-1β production. Careful analysis of our existing single-cell time course data does not allow us to distinguish between these two possibilities because modest elevations of IL-1β in macrophages and myeloid chemokines in fibroblasts are already detectable at 6 weeks of age. Novel “sensors” of circuit activation in the TME (such as IL-1β reporter macrophages or IL-6 reporter fibroblasts) may be required to identify these very early signals from tumor cells.

In future work it will also be important to clarify whether and how T cells participate in the IL-1β -> IL-6 TME circuit. Together with macrophages, T cells are a prominent feature of the immune cell infiltrate in both the Hi-Myc and PM models, with evidence of dysfunction based on expression of canonical exhaustion markers. Intriguingly, the exhaustion phenotype is reversed by macrophage depletion (consistent with myeloid-derived immune suppression), yet, based on CD8^+^ T cell depletion studies (**Fig 2N-O**), restoration of T cell function does not appear to play a role in the impaired invasion seen following macrophage depletion. Importantly, T cells have been documented to play a role in immune surveillance in prostate cancer in other contexts,^38,90^ but presumably independent of the IL-1β -> IL-6 TME circuit described here.

The fact that combined IL-1β+IL-6 blockade is superior to IL-1β blockade alone in our prostate cancer initiation model raises the question of whether clinical translation is warranted, particularly since this could be tested using inhibitors already approved for other indications.

However, it is important to acknowledge that prior clinical efforts targeting these or related pathways in prostate cancer have failed–but under a very different clinical context than proposed here. Specifically, a phase 2 clinical trial of the IL-6 neutralizing antibody siltuximab in metastatic castration resistant prostate cancer (mCRPC) patients showed minimal activity.^91^ Similarly, a phase 1 clinical trial of a CSF1R blocking antibody (targeting macrophages) had modest clinical benefit in only 3 of 12 patients with mCRPC and was not pursued further.^92^ These results, coupled with the lack of clinical response to IL-1β-neutralizing antibody (canakinumab) in advanced lung cancer,^47^ suggest that combined IL-1β + IL-6 blockade would have a low probability of success in late-stage prostate cancer. However, neoadjuvant therapy has been recently validated as a new treatment paradigm in men with high-risk localized prostate cancer based on phase 3 data showing clinical benefit of androgen deprivation therapy (ADT) plus an androgen receptor signaling inhibitor (apalutamide) prior to surgery.^93^ This clinical context could be an optimal setting for adding IL-1β + IL-6 blockade (assuming acceptable safety) because the pathologic response rate with ADT + apalutamide alone was only 9%, offering plenty of room for improvement.

Although preliminary, the potential relevance of the IL-1β -> IL-6 circuit in lung cancer is worth exploring based on recent work linking IL-1β to clonal hematopoiesis (CH), and to worse clinical outcomes in patients whose tumors are infiltrated with IL-1β^+^ macrophages derived from the CH clone.^94,95^ These findings, together with the reduction in new lung cancer cases in patients treated with IL-1β-neutralizing antibody in the CANTOS trial,^86^ have sparked a reevaluation of IL-1β blockade in lung cancer, but now in the context of lung cancer prevention using biomarker-based patient selection.^96^ If an IL-1β -> IL-6 circuit is confirmed in lung cancer, addition of IL-6 inhibition to such a prevention regimen is worth consideration, again assuming adequate safety.

### Limitations

We intentionally selected the Hi-Myc prostate cancer model for our study because it temporally separates the preneoplastic stage of tumor initiation from the stage of tissue invasion. The fact that the transcriptome of *TG-MYC*^+^ cells during the invasion stage is fully present at the PIN stage provided critical insight that changes in the TME are required before the full invasive program becomes manifest. That said, it is possible that additional cancer cell-intrinsic events also play a role. Indeed, in early work we documented androgen receptor gene amplification in a cell line derived from a Hi-Myc tumor, but this is a rare event.^97^

Hi-Myc mice serve as an ideal platform to study the PIN-to-PCa transition but are cumbersome for evaluating functional perturbations of the TME due to the large number of genetic crosses involved. To overcome this barrier, we developed the PM orthograft model which, importantly, reproduces the critical TME features seen in Hi-Myc mice such as infiltration of IL-1β^+^ macrophages and induction of IL-6 in prostate fibroblasts. The speed of the orthograft model provides the additional advantage of quickly evaluating multiple candidate TME perturbations, but it does not allow a clean separation of the PIN stage from the invasive cancer stage.

Our functional studies documenting the importance of the IL-1β -> IL-6 circuit have focused exclusively on MYC-driven prostate cancer models. We use transcriptomic data to provide evidence for this circuit in other prostate cancer models, and in human prostate and lung adenocarcinoma, but it will be important to confirm these findings at a functional level. Finally, our studies have, for obvious reasons, focused on IL-1β production in tumor-infiltrating macrophages. It is worth noting that IL-1α also plays a critical role in certain epithelial cancers, such as pancreatic cancer, in which IL-1α secreted by tumor cells reprograms normal pancreatic fibroblasts into inflammatory cancer-associated fibroblasts (iCAFs).^98^ Because IL-1α and IL-1β share a common receptor (IL-1R1), these and related findings provide a rationale for future therapeutic strategies targeting IL-1R1 rather than IL-1β alone.

## Acknowledgements

We thank members of the Sawyers laboratory for valuable advice and discussions, with special thanks to Zhendong Cao, Pan Cheng, and Teng Han. We are grateful to Rona Lester for help in mouse colony management, the Molecular Cytology Core Facility at MSKCC for help with microscopy, the Single Cell Analytics Innovation Lab Core Facility at MSKCC for help with single-cell library preparation, the Integrated Genomics Operation Core Facility at MSKCC for sequencing, the Flow Cytometry Core Facility at MSKCC for help with flow cytometry and FACS experiments, and the Antitumor Assessment Core Facility at MSKCC for *in vivo* studies. C.L.S. was supported by the Howard Hughes Medical Institute, Calico Life Sciences LLC, and NIH grants CA193837, CA092629, CA265768, CA008748. Y.S.L. was supported by the Barbara and Stephen Friedman Pre-doctoral Fellowship and a Medical Scientist Training Program grant from the National Institute of General Medical Sciences of the National Institutes of Health under award number: T32GM152349 to the Weill Cornell/Rockefeller/Sloan Kettering Tri-Institutional MD-PhD Program.

## Author Contributions

C.L.S., J.Z., and Y.S.L. conceived the project. C.L.S., J.Z., Y.S.L., A.C., M.L., B.S.C., D.P., and R.R. oversaw the project, performed experimental design and data interpretation. C.L.S., J.Z., and Y.S.L. co-wrote the manuscript. B.S.C., R.R., and A.C. edited the manuscript. M.L., Y.S.L., R.S., J.C., S.K., L.F., and Z.C. performed computational analysis. J.Z., O.C., T.X., I.M., and R.C. performed scRNA-seq experiments. J.Z., Y.S.L., A.C., P.S., and K.L. analyzed tumors and performed flow cytometry. R.R. provided reagents for PM tumoroids and cloning experiments. E.B. provided feedback throughout the project and protocol and reagents for *ex vivo* fibroblast experiments. H.Z., Y.S.L., J.Z., A.C., P.S., K.L., and E.S. oversaw and performed *in vivo* experiments. Y.S.L. performed cloning and Western blot experiments, conditioned media assays, fibroblast *ex vivo* studies, and qRT-PCR analysis. M.H. and A.G. reviewed and analyzed tissue slides. W.K., S.N., E.R., and N.F. performed tissue embedding, sectioning, slide scanning, and staining. All authors approved of the final manuscript.

---

**Supplemental Figure 1.**
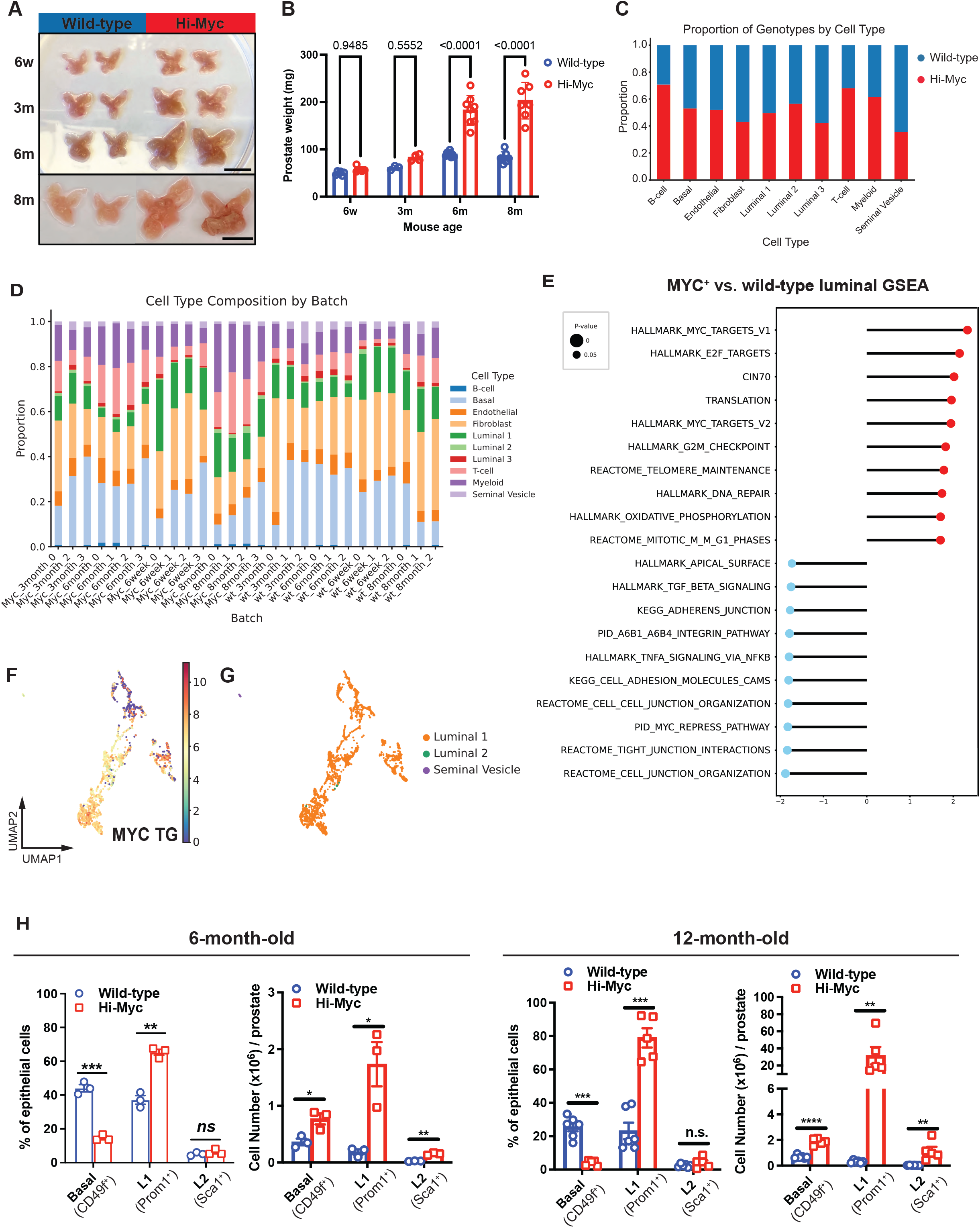
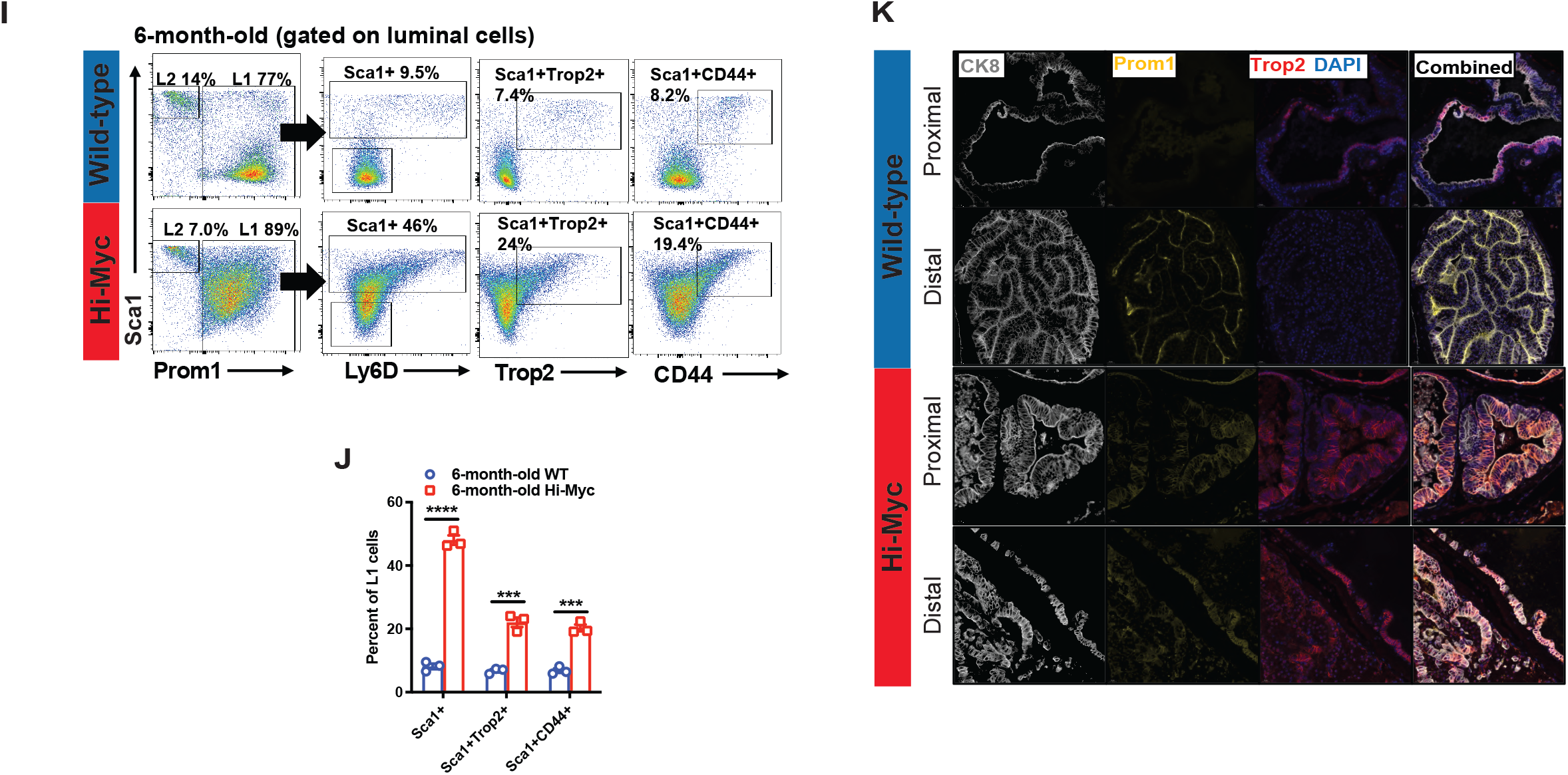
(A and B) Representative images (A) and prostate weight quantification (B) of age-matched wild-type and Hi-Myc mice. Scale bar: 10 mm. Unpaired two-tailed t-test; data are represented as mean ± s.d. (C) Stacked bar graph showing the proportion of genotypes across cell types in the scRNA-seq dataset. (D) Stacked bar graph showing the proportion of each cell type across individual samples (batches) in the scRNA-seq dataset, stratified by genotype and age. (E) Gene set enrichment analysis (GSEA) of MYC^+^ cells from Hi-Myc prostates compared with luminal epithelial cells from wild-type prostates. (F) UMAP visualization of transgene human *MYC* expression in Hi-Myc epithelial cells. (G) UMAP showing epithelial cell subtypes within Hi-Myc epithelial cells. (H) Quantification of Basal, Luminal 1, and Luminal 2 epithelial cells in Hi-Myc prostates from 6-month-old (left) and 12-month-old (right) mice as determined by flow cytometry. Unpaired two-tailed t-test; data are represented as mean ± s.d. (I and J) Flow cytometry gating strategy (I) and quantification (J) to identify Luminal 1 cells expressing stem cell-associated markers. Unpaired two-tailed t-test; data are represented as mean ± s.d. (K) Representative IF images of wild-type and Hi-Myc prostate tissues. Scale bar: 20 µm.

**Supplemental Figure 2.**
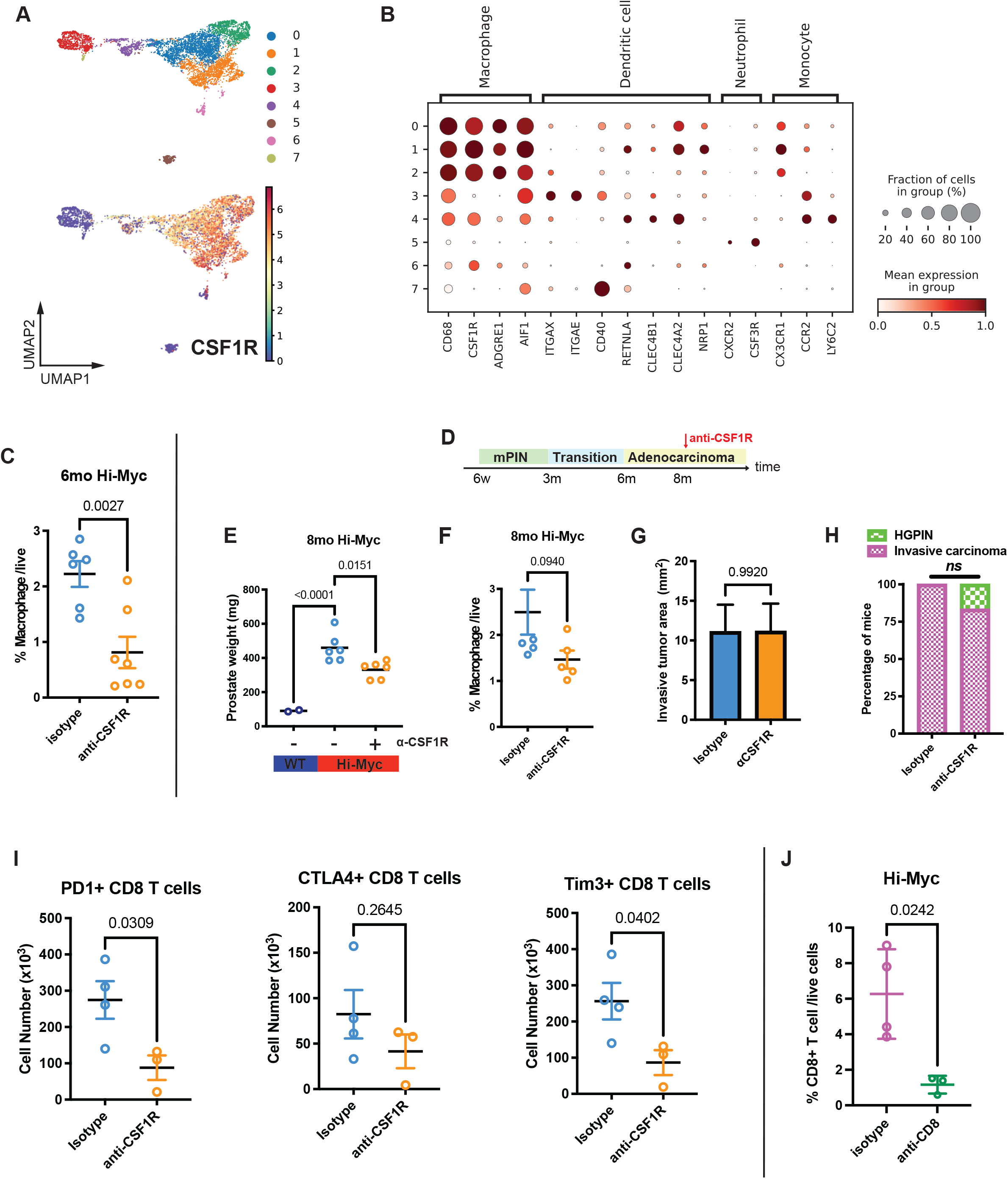
(A) UMAP visualization of Leiden clusters (top) and *Csf1r* gene expression (bottom) gene across wild-type and Hi-Myc myeloid populations. (B) Dot plot showing canonical myeloid cell markers used for cell type annotation in the scRNA-seq dataset. (C) Quantification of macrophages (CD45^+^ CD11b^+^ Ly6C^low^ Ly6G^-^ F4/80^+^) following antibody treatment of 6-month-old Hi-Myc mice. Unpaired two-tailed t-test; data are represented as mean ± s.d. (D) Schematic of a 10-week anti-CSF1R antibody treatment regimen in Hi-Myc mice, initiated at 8 months of age. 6w, 6-week-old; 3m, 3-month-old; 6m, 6-month-old; 8m, 8-month-old. (E) Prostate weights following antibody treatment of 8-month-old Hi-Myc mice as outlined in (D). One-way ANOVA; horizontal lines represent the mean. (F) Quantification of tumor infiltrating macrophages (CD45^+^ CD11b^+^ Ly6C^low^ Ly6G^-^ F4/80^+^) following antibody treatment of 8-month-old Hi-Myc mice. Unpaired two-tailed t-test; data are represented as mean ± s.d. (G and H) Quantification of invasive tumor area (G) and pathological tumor stage assessment performed by an external clinical pathologist (H) following anti-CSF1R antibody treatment of 8-month-old Hi-Myc mice. Unpaired two-tailed t-test; data are represented as mean ± s.d. (I) Quantification of indicated exhaustion marker expression in CD8^+^ T cells following acute 1-week antibody treatment. Unpaired two-tailed t-test; data are represented as mean ± s.d. (J) Quantification of CD8^+^ T cells (CD45^+^ CD3^+^ CD8^+^) in peripheral blood following acute 1-week antibody treatment. Unpaired two-tailed t-test; data are represented as mean ± s.d.

**Supplemental Figure 3.**
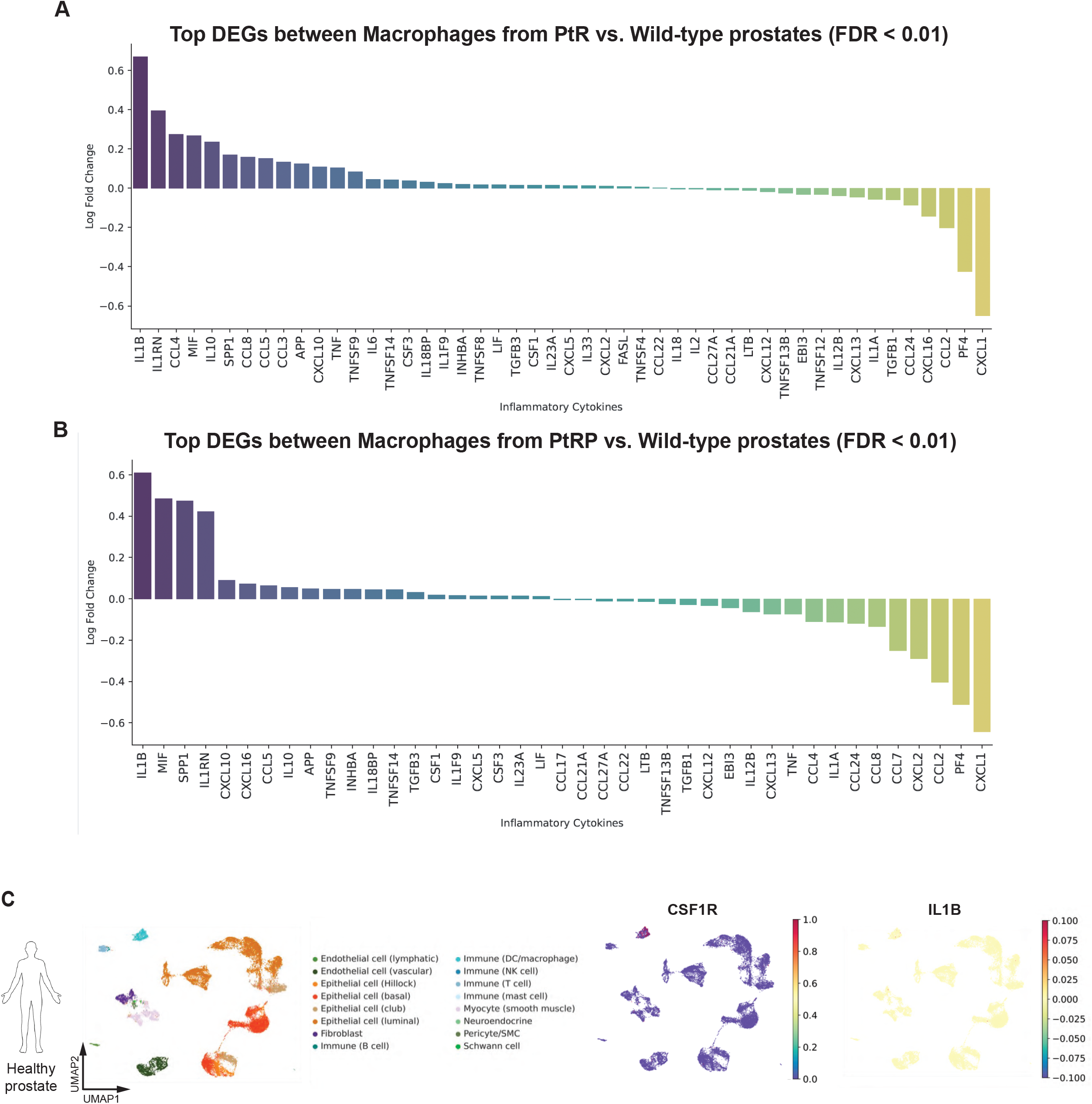
(A) Differentially expressed inflammatory cytokines in macrophages from PtR tumors compared with those from wild-type prostates (FDR<0.01). (B) Differentially expressed inflammatory cytokines in macrophages from PtRP tumors compared with those from wild-type prostates (FDR<0.01). (C) UMAP visualization of *CSF1R* and *IL1B* expression across cell types in healthy human prostate tissue. Single-nucleus RNA-sequencing (snRNA-seq) data were obtained from the Genotype-Tissue Expression (GTEx) Consortium cell atlas.

**Supplemental Figure 4.**
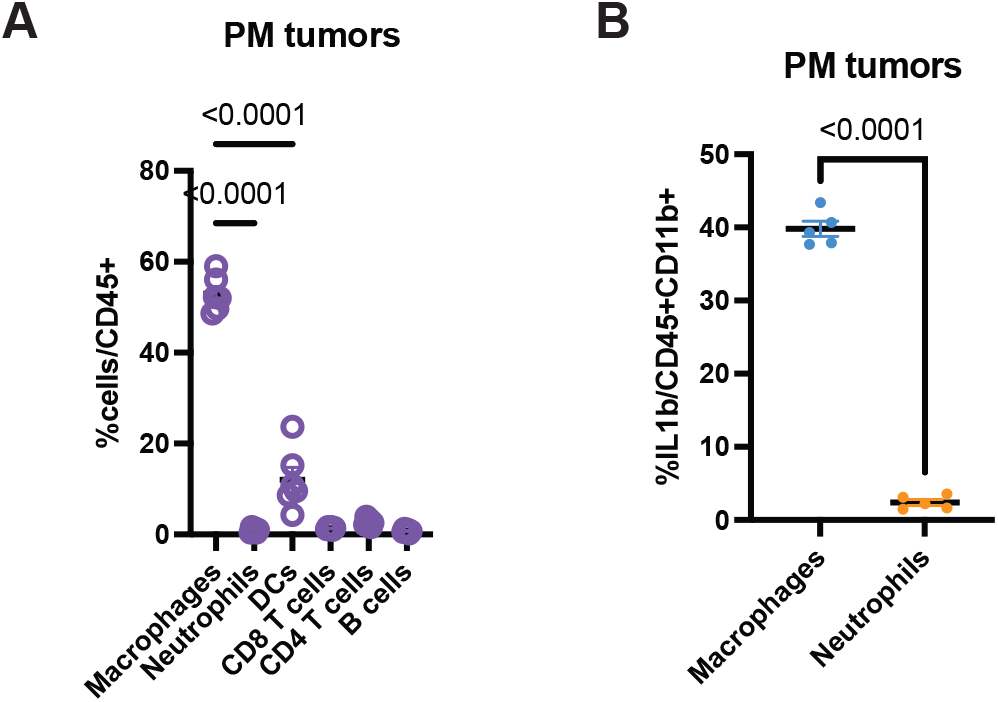
(A) Quantification of myeloid cell subsets in PM tumors, as determined by flow cytometry. One-way ANOVA; horizontal lines represent the mean. (B) Quantification of IL-1β^+^ cells in macrophages (CD45^+^ CD11b^+^ Ly6C^low^ Ly6G^-^ F4/80^+^) and neutrophils (CD45^+^ CD11b^+^ Ly6C^-^ Ly6G^+^) in PM tumors. Unpaired two-tailed t-test; data are represented as mean ± s.d.

**Supplemental Figure 5.**
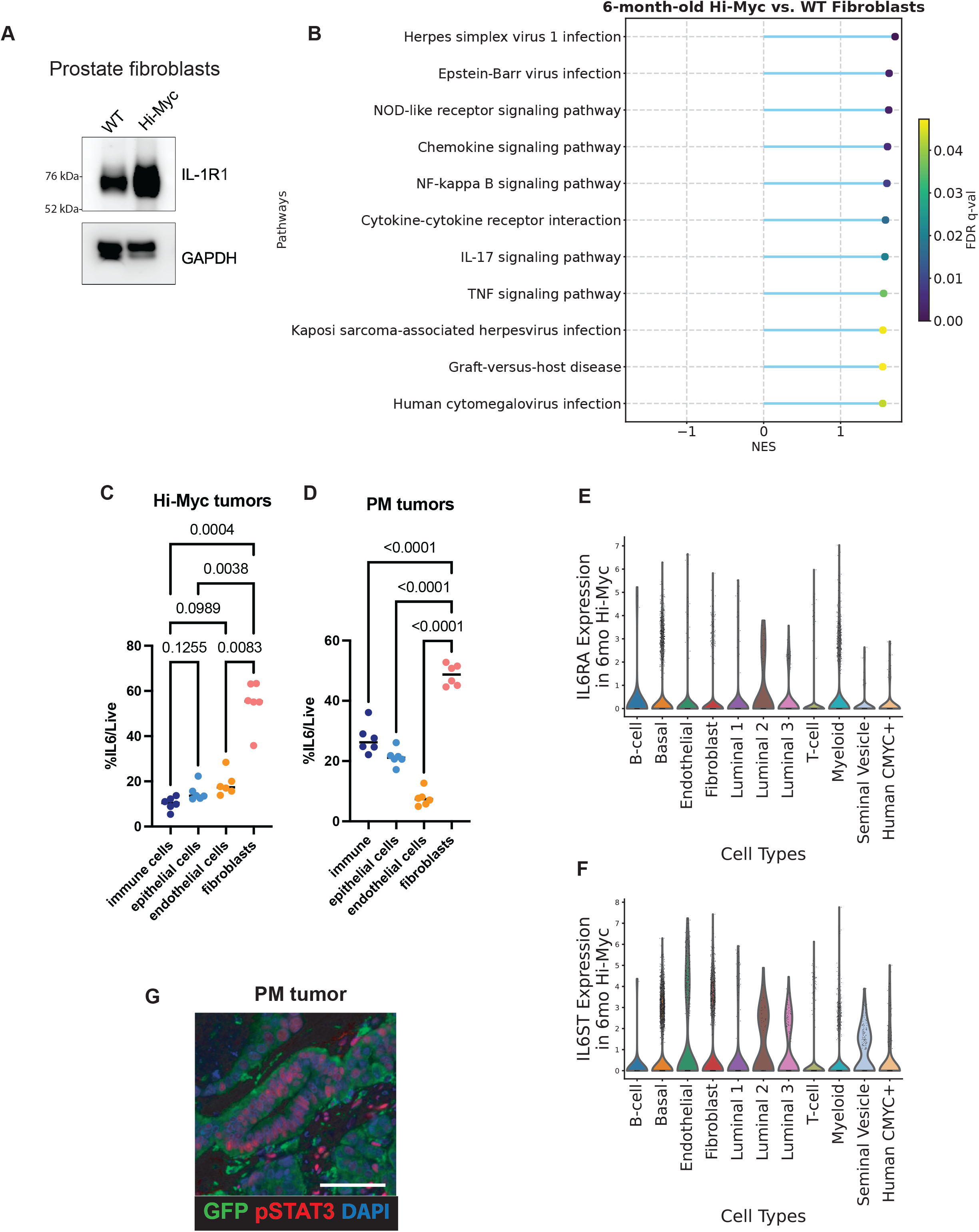
(A) Western blot showing IL-1R1 levels in prostate fibroblasts from 6-month-old Hi-Myc or age-matched wild-type mice. GAPDH was used as a loading control. (B) Dot plot showing GSEA pathways enriched in fibroblasts from 6-month-old Hi-Myc prostates compared with fibroblasts from age-matched wild-type prostates. (C and D) Quantification of IL-6^+^ cells in immune cells (CD45^+^), epithelial cells (CD45^-^ EpCAM^+^), endothelial cells (CD45^-^ EpCAM^-^ CD31^+^), and fibroblasts (CD45^-^ EpCAM^-^ CD31^-^ PDGFRα^+^) in 6-month-old Hi-Myc tumors (C) and PM tumors (D). One-way ANOVA; horizontal lines represent the mean. (E and F) Violin plots showing *Il6ra* (E) and *Il6st* (F) gene expression across cell types in 6-month-old Hi-Myc prostates. (G) Representative multiplex immunofluorescence image of phospho-STAT3^+^ cancer cells (GFP^+^) in PM tumors. Scale bar: 50 µm.

**Supplemental Figure 6.**
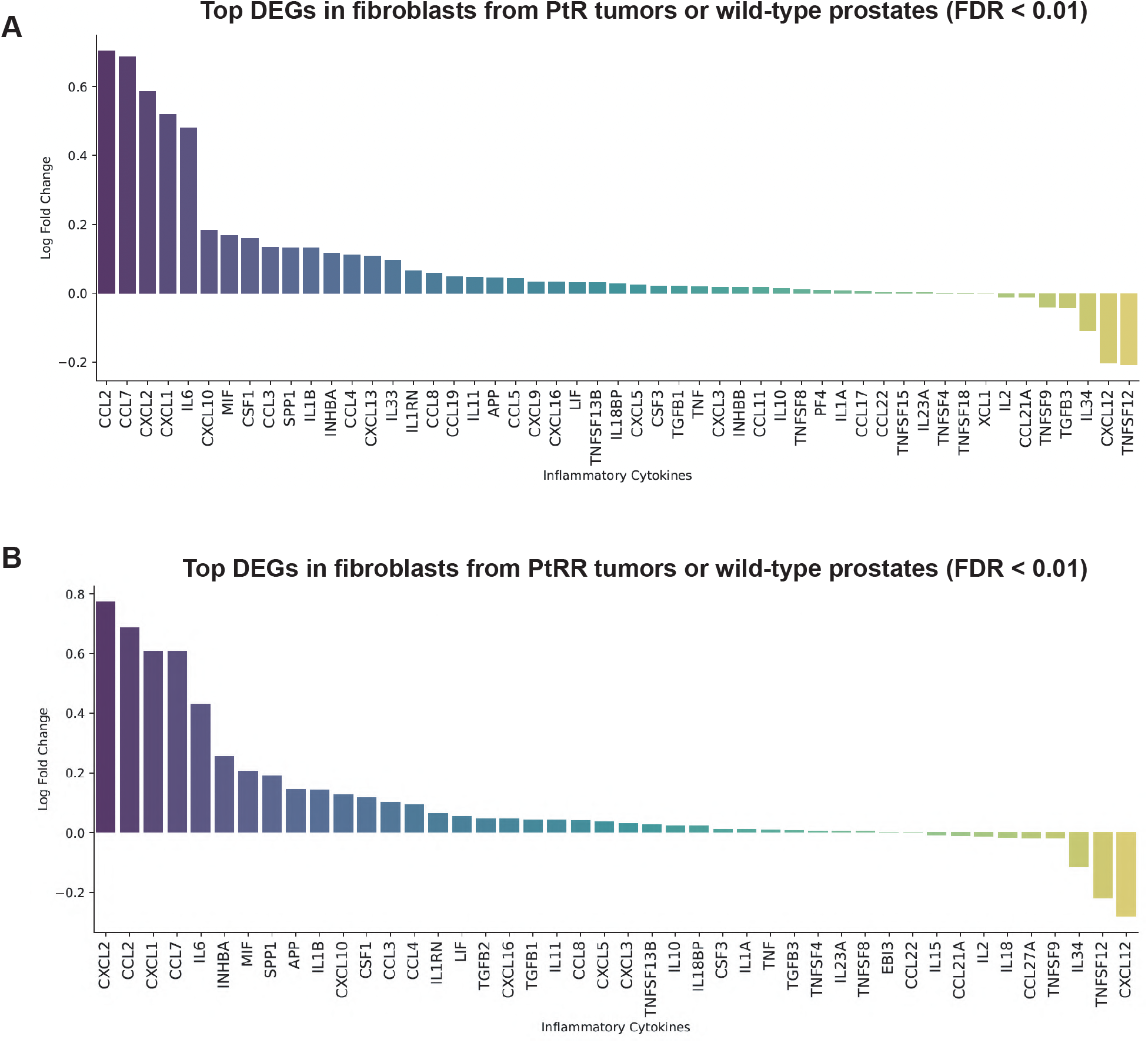
(A) Differentially expressed inflammatory cytokines in fibroblasts from PtR tumors compared with those from wild-type prostates (FDR<0.01). (B) Differentially expressed inflammatory cytokines in fibroblasts from PtRP tumors compared with those from wild-type prostates (FDR<0.01).

## METHODS

### Mouse models

Hi-Myc mice generated in our laboratory were used in this study.^48^ C57BL/6J (JAX:000664) and B6.129S2-Il6^tm1Kopf^/J (JAX:002650) mice were purchased from The Jackson Laboratory.

B6.129S7-Il1r1^tm1Imx^/J (JAX:003245) mice were kindly provided by the Tobias Hohl laboratory (MSKCC). All mice were maintained on a C57BL/6 background. Upon arrival, mice were acclimated for at least 1 week prior to experimentation.

Male mice were used for all experiments. Hi-Myc mice ranged in age from 6 weeks to 8 months, with exact ages specified in the corresponding figure legends or Results section. All other mouse strains were used between 8 and 12 weeks of age. Prostate cancer is a male-specific disease; therefore, only male mice were included.

Mice were randomly assigned to experimental groups where applicable. All animal studies and procedures were approved by the Memorial Sloan Kettering Cancer Center Institutional Animal Care and Use Committee (IACUC protocol 06-07-012) and were conducted in accordance with institutional guidelines for the humane use of animals in biomedical research, including approved methods of euthanasia.

### Tumoroid cell line

The *Trp53^-/-^*; *Myc^T58A^* (PM) tumor-derived organoid line (tumoroid) was generated from an orthotopically-transplanted PM organoid model that was established as previously described.^10^ Briefly, following targeted *Trp53* deletion via Cas9 nucleoprotein electroporation and transduction of *Myc^T58A^-2A-EGFP* overexpression construct, organoids were orthotopically implanted into GFP-tolerant C57BL/6 mice (JAX#: 026175). Tumors were harvested, dissociated, and re-cultured to establish tumoroids.

### Public human datasets

Publicly available human transcriptomic datasets were used in this study. The Cancer Genome Atlas (TCGA) RNA-seq expression data and corresponding clinical annotations were obtained from cBioportal for Cancer Genomics.^99,100^ The results are in whole or part based upon data generated by the TCGA Research Network (https://www.cancer.gov/tcga/). Normal tissue RNA sequencing data were obtained from the Genotype-Tissue Expression (GTEx) portal (https://www.gtexportal.org).^101^ The GTEx Project was supported by the Common Fund of the Office of the Director of the National Institutes of Health, and by NCI, NHGRI, NHLBI, NIDA, NIMH, and NINDS.

Single-cell RNA sequencing (scRNA-seq) datasets were obtained from previously published studies, including Song *et al.*,^40^ Hirz *et al.*,^39^ and Lyu *et al*.^38^ Raw and processed gene expression matrices and associated metadata were downloaded from the Gene Expression Omnibus (GEO) repository under the accession numbers reported in the original publications.

All datasets were de-identified and publicly available and thus did not require additional institutional review board approval. Datasets were accessed in accordance with the guidelines of the respective repositories. Sample sizes, patient characteristics, and additional metadata are described in the original publications and summarized where relevant in the Results.

#### *In vivo* mouse studies

##### Prostatic orthotopic transplantation

8–12-week-old male mice were randomized for surgical implantation into a single dorsal prostatic lobe as described previously.^102^

##### Antibody treatment

Mice were treated with anti-CSF1R (300 μg/dose), anti-CD8a (300 μg/dose), anti-PD1 (200 μg/dose), anti–IL-1β (100 μg/dose), anti–IL-1R1 (500 μg/dose), or anti–IL-6R (500 μg/dose), or isotype control antibodies (details provided in the Key Resources Table). Antibodies were administered via intraperitoneal injection in a volume of 200 μL per dose, three times per week for the duration of the experiment. Treatment was continued for 10 weeks in Hi-Myc mice and 3 weeks in PM tumoroid-bearing mice.

##### Tissue collection

Mice were euthanized and tumors were excised and weighed prior to processing. For generation of single-cell suspensions, tumor tissues were minced and enzymatically digested in 5 mL of collagenase solution consisting of Collagenase type XI (0.4 mg/mL), 0.36 mM CaCl₂, and 20 μg/mL DNase I, at 37°C for 30 minutes. Digestion was neutralized with RPMI-based buffer supplemented with 5% heat-inactivated fetal bovine serum, GlutaMAX, penicillin–streptomycin, and 5 mM HEPES. The resulting cell suspension was filtered through a 70 μm cell strainer.

##### Histology and immunohistochemistry

Prostate tissues were fixed in 4% paraformaldehyde, paraffin embedded, and sectioned at 5 μm. Sections were subjected to hematoxylin and eosin (H&E) staining or chromogenic immunohistochemistry (IHC). For IHC, sections were deparaffinized and subjected to heat-induced antigen retrieval using Cell Conditioning 1 (Ventana). Primary antibodies (listed in the Key Resources Table) were detected using HRP-conjugated secondary reagents and visualized with DAB chromogen. Slides were counterstained with hematoxylin, dehydrated, and mounted. Whole-slide images were acquired using a Pannoramic Scanner (3DHistech) with a ×20/0.8 NA objective and analyzed using ImageJ or QuPath (v0.5.1).

##### Multiplexed immunofluorescence

Multiplexed immunofluorescence staining was performed using a Leica Bond automated staining platform (Leica Biosystems). Paraffin-embedded sections (5 μm) were subjected to antigen retrieval using EDTA-based ER2 solution (Leica, AR9640), followed by sequential rounds of primary antibody staining, HRP-conjugated secondary detection, and tyramide signal amplification. Antigen retrieval was repeated between staining cycles to denature previously bound antibodies prior to subsequent rounds of staining. Sections were counterstained with DAPI, rinsed in PBS, mounted, and imaged.

#### *Ex vivo* primary cell assays

##### Fibroblast isolation

Mouse prostates were dissected and dissociated as described above. Dissociated cells were plated without prior filtration onto tissue culture–treated 10 cm plates (one prostate per plate) in DMEM supplemented with 10% HI-FBS, penicillin–streptomycin, GlutaMAX, and 5 mM HEPES. Cells were washed with PBS and media was replaced the following day to remove non-adherent cells. After 5 days of culture, adherent cells were subjected to negative selection to enrich for fibroblasts.

Fibroblasts were isolated using the EasySep™ Mouse Streptavidin RapidSpheres™ Isolation Kit (Stemcell Technologies) according to the manufacturer’s instructions and as previously described (cite relevant studies). Briefly, cells were incubated with biotinylated antibodies against CD45, EpCAM, and CD31, followed by magnetic bead–based depletion to remove immune cells (CD45⁺), epithelial cells (EpCAM⁺), and endothelial cells (CD31⁺). The remaining negatively selected fibroblast population was used for downstream analyses. The efficacy of fibroblast enrichment was validated by flow cytometry. Primary fibroblasts were maintained in culture for a maximum of two passages.

#### *In vitro* cell-based assays

##### Gene editing (CRISPR RNP and lentiviral)

CRISPR ribonucleoprotein (RNP)–mediated gene editing was performed as previously described.^102^ Briefly, recombinant Cas9 protein was complexed with synthetic single guide RNAs (sgRNAs) targeting *Il1r1* or *Il6st*, designed using CRISPRick and purchased from Integrated DNA Technologies (IDT) (sequences provided in the Key Resources Table). RNP complexes were delivered into PM tumoroids via electroporation.

For *Il1r1*-knockout tumoroids generated via CRISPR RNP, loss of IL-1R1 expression was confirmed by immunoblotting. For *Il6st* knockout, cells were stained for gp130 surface expression, and the gp130-negative population was enriched by FACS.

For lentiviral-mediated gene editing, U6-sgRNA-EFS-mScarlet (USEmS) constructs were generated by Gibson assembly as previously described.^103^ PM tumoroids were transduced with lentiviral vectors encoding sgRNAs targeting *Il1r1* along with an mScarlet reporter. Transduced (mScarlet⁺) cells were enriched by FACS.

##### RT-qPCR

Fibroblasts from wild-type or Hi-Myc prostates were seeded at equal densities and allowed to adhere overnight prior to stimulation. Cells were treated with recombinant IL-1β for 4 hours, with 3 independent biological replicates per condition. Total RNA was extracted using the RNeasy kit (Qiagen) according to the manufacturer’s instructions. cDNA was synthesized using the High-Capacity cDNA Reverse Transcription Kit (Thermo Fisher Scientific).

Quantitative PCR was performed using PowerTrack SYBR Green Master Mix (Thermo Fisher Scientific) on a QuantStudio 7 Flex Real-Time PCR System (Applied Biosystems). Relative gene expression was calculated using the ΔΔCt method, with *Actb* (β-actin) as the internal control.

##### Cytokine quantification in conditioned media

Fibroblasts from wild-type or Hi-Myc prostates were seeded at equal densities and allowed to adhere overnight. Conditioned media was collected after 24 hours of culture. Media samples were centrifuged at 600 × g for 5 minutes to remove cellular debris, and the clarified supernatant was collected. Conditioned media samples were submitted to Eve Technologies for multiplex cytokine analysis using a Luminex platform. Six independent biological replicates were analyzed per condition.

##### Immunoblotting

Cell lysates were prepared in RIPA buffer and sonicated. Protein concentrations were determined using a BCA assay (Thermo Fisher Scientific). Equal amounts of protein were mixed with NuPAGE LDS sample buffer, denatured at 95°C, and resolved on 4–12% Bis–Tris gels (Thermo Fisher Scientific). Proteins were transferred to PVDF membranes (Millipore), blocked in 5% milk in TBST, and incubated with primary antibodies overnight at 4°C. Membranes were washed and incubated with HRP-conjugated secondary antibodies prior to detection using ECL substrate (Cytiva). Blots were imaged on an ImageQuant 800 system (Cytiva).

##### Flow cytometry

Prior to staining, a Percoll density gradient was used to enrich viable cells from Hi-Myc tumor samples. Briefly, cells were resuspended in 40% Percoll (diluted in HBSS) and layered over 80% Percoll to form a discontinuous gradient. Samples were centrifuged at 300 × g for 30 minutes at room temperature with low acceleration and deceleration. Debris in the upper phase was removed, and cells from the intermediate phase were collected, washed with HBSS, and pelleted by centrifugation at 600 × g for 5 minutes. Percoll enrichment was not performed for PM tumoroid-derived samples.

Following preparation of single-cell suspensions, cells were resuspended in flow cytometry buffer consisting of PBS supplemented with 2% heat-inactivated fetal bovine serum and 1 mM EDTA. Cells were incubated with Fc receptor–blocking antibody for 15 minutes at 4°C, washed, and stained with a live/dead viability dye (1:500 dilution in PBS) for 10 minutes at 4°C. Cells were washed again and incubated with fluorophore-conjugated primary antibodies diluted 1:200 in flow cytometry buffer for 15 minutes at 4°C (antibodies listed in the Key Resources Table).

For intracellular staining, cells were fixed using fixation buffer for 15 minutes at 4°C, followed by overnight incubation with secondary antibodies at 4°C. After staining, cells were washed, resuspended in flow cytometry buffer, and transferred to round-bottom 96-well plates for analysis.

Flow cytometry was performed using a CytoFLEX flow cytometer (Beckman Coulter). Data were analyzed using FlowJo v10.9.0 (BD Biosciences).

##### Fluorescence-activated cell sorting (FACS)

Cells were stained with DAPI (1 μg/mL in PBS) for 15 minutes at 4°C, washed in flow cytometry buffer (PBS supplemented with 2% HI-FBS and 1 mM EDTA), and filtered through a 35 μm nylon mesh to obtain a single-cell suspension. Target cell population was sorted using a BD FACSAria III cell sorter (BD Biosciences).

#### Single-cell RNA-seq analysis

##### Single-cell RNA-seq preprocessing, integration, and cell-type annotation

Raw scRNA-seq FASTQ files were processed using SEQC to generate gene-by-cell count matrices. Empty droplets were identified and removed using EmptyDrops. Genes detected in fewer than 10 cells were excluded from downstream analyses. Principal component analysis (PCA) was performed on library-size normalized expression values and used for quality-control clustering. Cells were clustered using PhenoGraph (k = 30), and clusters exhibiting elevated mitochondrial transcript content were removed. Following quality-control filtering, 67,776 cells and 18,278 genes were retained for downstream analyses.

Batch effects were assessed using neighborhood batch entropy calculated from replicate labels. For each cell, replicate frequencies among the k = 30 nearest neighbors were converted into a probability distribution and Shannon entropy was calculated. Batch correction was performed using Scanorama (k = 20, dimred = 100, alpha = 0.1) in a two-step procedure. Replicates were first integrated within each timepoint, followed by integration across timepoints. UMAP embeddings were generated from PCA-reduced and Scanorama-corrected expression matrices. Highly variable genes were identified using the Scanpy pp.highly_variable_genes function. Cells were clustered using PhenoGraph on PCA-reduced expression matrices. Broad cell identities, including epithelial, immune, mesenchymal, and endothelial populations, were assigned using canonical marker genes and cluster-level expression profiles. Differentially expressed genes were identified using MAST on log-normalized expression matrices with default parameters. Unless otherwise indicated, each cluster was compared against all remaining cells. Fine cell-state annotations were assigned using canonical markers together with MAST-derived differentially expressed genes.

##### Epithelial cell-state analysis

Wild-type epithelial cells were isolated and reclustered using PhenoGraph. Cell identities were assigned using canonical epithelial markers and MAST-derived differentially expressed genes. Cell-state labels were subsequently transferred to Hi-Myc epithelial cells using the PhenoGraph classifier on Scanorama-corrected expression matrices. A curated panel of 468 epithelial lineage marker genes was used as classifier features. MYC-positive tumor cells were identified using MAGIC-imputed HUMANCMYC expression (k = 30, t = 3). Cells with imputed HUMANCMYC expression greater than 2 were classified as MYC-positive.

Differential expression analyses between epithelial populations were performed using MAST. Genes were ranked according to signed −log10(FDR) values and analyzed using preranked GSEA implemented in GSEApy.

Latent transcriptional programs were identified using single-cell hierarchical Poisson factorization (scHPF). scHPF was performed on raw count matrices with 10 latent factors, and factor activities were compared across tumor progression timepoints.

##### Myeloid and fibroblast analyses

Myeloid and fibroblast populations were isolated from the integrated dataset and analyzed independently. Highly variable genes were identified, principal component analysis was performed, and UMAP embeddings were generated for visualization. Clustering was performed using the Leiden algorithm. Cluster identities were assigned using canonical lineage-specific marker genes together with cluster-level expression profiles.

Tumor-infiltrating macrophages were defined as macrophages present in Hi-Myc tumors. Differentially expressed genes between Hi-Myc tumor-infiltrating and corresponding wild-type populations were identified using MAST with default parameters. Pathway enrichment analyses were performed using the same GSEApy-based workflow described above.

##### TCGA transcriptomic analysis

TCGA Firehose Legacy RNA-seq expression data and corresponding clinical annotations were obtained through cBioPortal for Cancer Genomics. Hi-Myc tumor-infiltrating macrophage and fibroblast gene signature scores were generated from genes upregulated in Hi-Myc tumor-infiltrating macrophages and fibroblasts relative to their corresponding age-matched control populations. Differentially expressed genes with log2 fold change ≥ 0.5 and FDR ≤ 1 × 10^-4^ were ranked by effect size, mapped to human orthologs, and used for gene signature score construction.

Gene signature scores were calculated using the Scanpy score_genes function. For TCGA Prostate Adenocarcinoma (PRAD), patients were stratified into low-risk (Gleason score ≤ 3+4) and high-risk (Gleason score ≥ 4+3) groups. Differences in macrophage and fibroblast gene signature scores between groups were evaluated using two-sided Mann–Whitney U tests.

Across cancer types, macrophage and fibroblast gene signature scores were calculated from TCGA bulk RNA-seq datasets and compared using Spearman correlation analysis.

## QUANTIFICATION AND STATISTICAL ANALYSIS

### Statistics and reproducibility

Statistical analyses were performed using GraphPad Prism (v10.5.0) and Python. Statistical tests used for individual experiments are indicated in the corresponding figure legends. Comparisons between two groups were performed using two-tailed Student’s t-tests or Mann– Whitney U tests, as appropriate. Multiple-group comparisons were performed using one-way or two-way ANOVA. Correlations were assessed using Spearman correlation analysis. Differential gene expression analyses were performed using MAST. Data are presented as mean ± SD unless otherwise indicated. Statistical significance was defined as ****p < 0.0001, ***p < 0.001, **p < 0.01, and *p < 0.05. Blinding was applied during group allocation, data collection, and analysis.

### Data availability

The data generated in this study as well as study resources are available upon request from the corresponding author.

